# ATRX Loss Predicts Poor Outcomes and Reveals a Therapeutic Vulnerability to TEAD Inhibition in Soft Tissue Sarcomas

**DOI:** 10.1101/2025.11.23.689992

**Authors:** Ryan A. Denu, Anand K. Singh, Yingda Jiang, Zhao Zheng, Davis R. Ingram, Khalida M. Wani, Rossana Lazcano, Madison P. Ginn, Kumaren Anand, Bishal K. Singh, Jackson C. Ballard, Veena Kochat, Biji Chatterjee, William Padron, Sharon M. Landers, Angela D. Bhalla, Asha Multani, Sharvari Dharmaiah, Prit Benny Malgulwar, Yiming Cai, Heather Lin, Jason T. Huse, Haoqiang Ying, Emily Z. Keung, Wei-Lien Wang, Anthony P. Conley, Shreyaskumar Patel, Neeta Somaiah, Ahsan Farooqi, Alexander J. Lazar, Elise F. Nassif Haddad, Keila E. Torres, Kunal Rai

**Affiliations:** Department of Sarcoma Medical Oncology, The University of Texas MD Anderson Cancer Center, Houston, TX; Department of Genomic Medicine, Division of Cancer Medicine, The University of Texas MD Anderson Cancer Center, Houston, TX; Department of Translational Molecular Pathology, The University of Texas MD Anderson Cancer Center, Houston, TX; University of Texas Health Sciences Center Houston, Houston, TX; Rice University, Houston, TX; Department of Surgical Oncology, The University of Texas MD Anderson Cancer Center, Houston, TX; Department of Genetics, The University of Texas MD Anderson Cancer Center, Houston, TX; Department of Molecular and Cellular Oncology, The University of Texas MD Anderson Cancer Center, Houston, TX; Department of Biostatistics, The University of Texas MD Anderson Cancer Center, Houston, TX; Department of Radiation Oncology, The University of Texas MD Anderson Cancer Center, Houston, TX; Department of Investigational Cancer Therapeutics, The University of Texas MD Anderson Cancer Center, Houston, TX

**Author notes:** Corresponding authors: Kunal Rai, The University of Texas MD Anderson Cancer Center, Houston, TX 77030., Ryan Denu, The University of Texas MD Anderson Cancer Center, Houston, TX 77030. Contributed equally (Anand K. Singh, Yingda Jiang, Zhao Zheng).

**Keywords:** ATRX, sarcoma, genomic instability, PRDM4, NFIX, Hippo pathway, YAP, TEAD, ALT, telomere

## Abstract

*ATRX* is one of the most frequently altered genes in sarcoma and encodes an ATP-dependent chromatin remodeler implicated in maintaining heterochromatin. However, *ATRX* alterations have not been leveraged for sarcoma treatment. We observed loss of ATRX protein in 14% of soft tissue leiomyosarcoma (STLMS, n =127), 53% of uterine leiomyosarcoma (ULMS, n = 95), 37% of undifferentiated pleomorphic sarcoma (UPS, n = 82), and 8% of dedifferentiated liposarcoma (DDLPS, n = 84). ATRX loss was associated with significantly worse outcomes in ULMS, UPS, and DDLPS. *ATRX* knockout in sarcoma cells increased proliferation in cooperation with *TP53* deletion. *ATRX* knockout led to chromatin de-repression and enrichment of *PRDM4* and *NFIX* transcription factor (TF) motifs. *PRDM4* and *NFIX* knockdown in *ATRX*-mutant sarcoma lines resulted in reduced proliferation and invasion suggesting epistatic relationship. Consistent with the known functional relationship between PRMD4 and YAP1, we observed that *ATRX/TP53* KO cells were more sensitive to the TEAD inhibitor VT103 compared to *TP53* KO and *ATRX* WT controls. Overall, our results identify *ATRX* loss as a prognostic factor of worse outcomes, implicate the ATRX-PRDM4-YAP1 axis as a novel underlying mechanism, and suggest use of TEAD inhibition as a potential therapeutic strategy for *ATRX*-deficient sarcomas.

**GRAPHICAL ABSTRACT:** 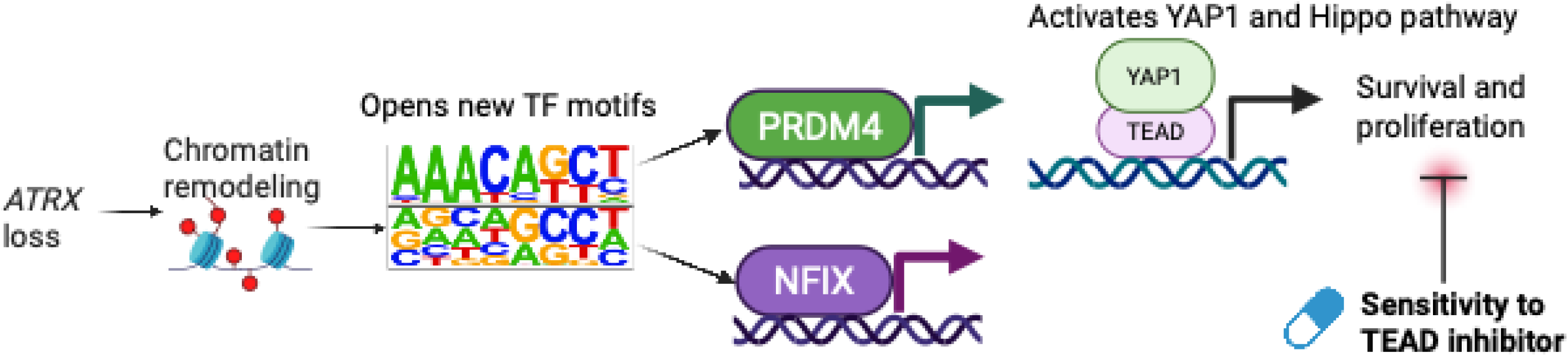

## INTRODUCTION

Sarcomas are heterogeneous mesenchymal malignancies with poor prognosis and limited treatment options in the metastatic setting. Treatment of recurrent or metastatic sarcoma typically involves combination cytotoxic chemotherapy for which response rates are rather poor (∼10-30%) and toxicity is high (1). There are limited predictive biomarkers available to help us know who is most likely to benefit from these aggressive chemotherapy regimens. In addition to optimizing our currently available therapies, there is a great unmet need to better understand the biology of sarcomas and to develop novel and more effective therapies.

Improved understanding of the cancer genome has led to many successes across oncology, especially with targeted therapies. However, this has not been the reality for many subtypes of sarcoma, as clinically targetable alterations are rarely seen. Some sarcoma subtypes are driven by fusion oncogenes, while others have complex genomes with prevalent gains and losses of chromosomes or chromosome arms, and point mutations driving oncogene overactivity are rarely seen. One of the most frequently altered genes in sarcoma is *ATRX*, which encodes an ATP-dependent chromatin remodeler with many different functions including epigenetic regulation, response to DNA damage, and telomere maintenance (2–5). *ATRX* was first discovered while discerning the cause of a syndrome characterized by anemia and intellectual disability, now called alpha thalassemia X-linked intellectual disability syndrome (6). *ATRX* is enriched in repetitive and GC-rich sequences in the genome and, in cooperation with the histone chaperone *DAXX*, is involved in depositing histone H3 variant 3.3 (H3.3) into the genome at areas of heterochromatin and pericentromeric regions, which helps to maintain these chromatin regions in a compact or silenced state (7, 8). Loss of *ATRX* increases G-quadruplexes, which are secondary DNA structures found at GC-rich regions of the DNA and are implicated in transcriptional dysregulation and DNA damage, as the presence of G-quadruplexes results in stalled replication forks and DNA damage (9). Concordantly, ATRX binding was enriched at G4 consensus motifs (10).

*ATRX* alterations are seen most commonly in sarcomas with complex genomes, such as uterine leiomyosarcoma (ULMS), in which loss-of-function mutations are seen in ∼30% (11–14). *ATRX* is also seen mutated in high-grade gliomas (15), pancreatic neuroendocrine tumors (16, 17), and neuroblastoma (18) but rarely in other cancer types, suggesting that the effects of *ATRX* loss are different based on the cell/tissue of origin. *ATRX* mutation is associated with worse outcomes in pancreatic neuroendocrine tumors (19) but better outcomes in gliomas (20, 21). In the soft tissue sarcoma (STS) TCGA dataset, *ATRX* mutation was associated with worse disease-specific survival in only the cohort of patients that did not receive radiation (22). However, the clinical significance of *ATRX* mutation is not well defined in different sarcoma subtypes. Further, the underlying mechanisms of *ATRX* alteration-mediated pathogenesis and effective treatments in this subgroup of sarcoma have not been identified.

## RESULTS

### ATRX mutation is common in specific sarcoma subtypes and co-occurs with TP53 alterations

*ATRX*, a known epigenetic regulator, is among the most commonly mutated genes in sarcoma (**Supplemental Figure 1**). Examination of different sarcoma subtypes in the AACR GENIE database (23, 24) showed that *ATRX* alterations were most frequent in ULMS (33% of ULMS with *ATRX* alteration), pleomorphic liposarcoma (PLPS, 21%), STMLS (20%), and UPS (20%) (**Figure 1A**). Similarly, in TCGA sarcoma dataset (25), *ATRX* alterations were most commonly seen in UPS (29.4%), LMS (14.3%), and DDLPS (10.7%; **Figure 1B**), and *ATRX* alteration was associated with worse overall survival (OS) (**Supplemental Figure 2**), as has been shown previously (22).

**Figure 1.**
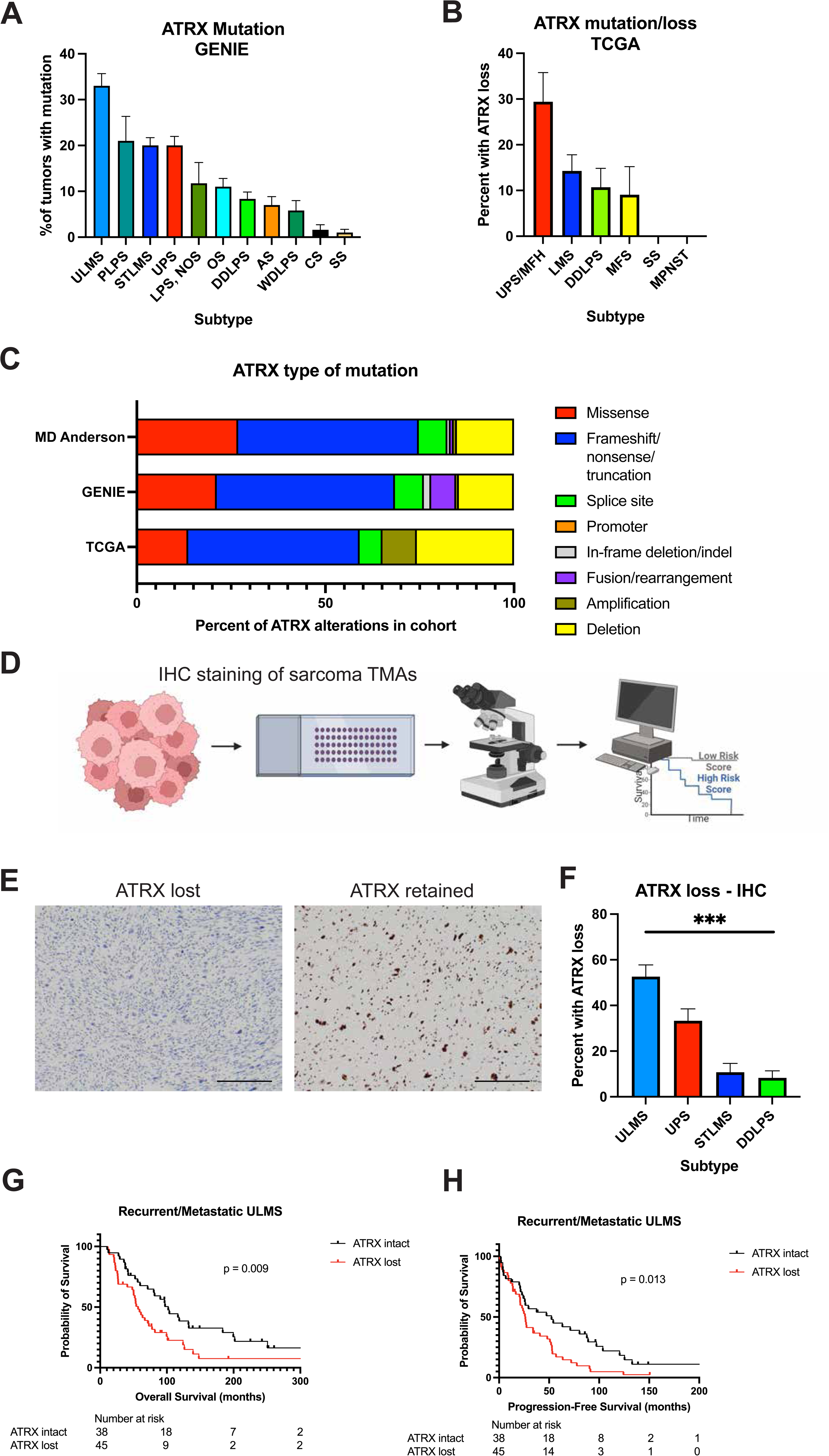
ATRX loss is associated with worse outcomes in multiple sarcoma subtypes. (A-B) Bar graph showing percent of each sarcoma subtype with *ATRX* mutation in the AACR GENIE database (A) and the TCGA database (B). In A-B, error bars represent standard error about the mean. (C) Distribution of *ATRX* alterations in our MD Anderson cohort, GENIE database, and TCGA database. Each color represents a different type of alteration. (D) Schemic of assessment of ATRX loss in sarcoma TMAs. (E) Representative images of loss of ATRX protein expression versus retention of ATRX expression. Scale bars = 100 μm. (F) Percent of tumors within each subtype with loss of ATRX by IHC. (G-H) Kaplan Meier curves displaying overall survival (G) and progression-free survival (H) in the recurrent/metastatic ULMS cases. P values displayed on Kaplan Meier curves are from log-rank tests.

We asked whether ATRX loss is prognostic in different sarcoma subtypes in additional cohorts of patients. As most *ATRX* alterations are deletions, frameshift or nonsense mutations (**Figure 1C**), they are predicted not to produce functional protein (26). We performed ATRX immunohistochemistry (IHC) in tissue microarrays (TMAs) containing 126 soft tissue LMS, 95 ULMS, 84 UPS, and 84 DDLPS (**Table S1; Figure 1D-E**). ATRX loss was seen in 52.6% of ULMS, 33.3% of UPS, 14.3% of STLMS, and 8.3% of DDLPS (**Figure 1F, Table S2**). We stratified survival analyses based on whether patients presented with localized or metastatic disease. In the ULMS cohort, 50/95 tumors demonstrated loss of ATRX. ATRX loss was associated with worse overall survival (OS) and progression free survival (PFS) in the metastatic setting in ULMS (**Figure 1G-H**). This was also seen in multivariate analysis (**Table S3**). In advanced/metastatic UPS, ATRX loss was associated with worse OS but no difference in PFS (**Supplemental Figure 3**). In localized UPS, there was no association between ATRX loss and RFS. In primary/localized STLMS, there was no significant difference in RFS (p=0.09), though this analysis is limited by small sample size (n = 66, 7 with ATRX loss and 59 with retained ATRX). Lastly, in metastatic DDLPS, ATRX loss was associated with significantly shorter PFS (p = 0.02) but no significant difference in OS (**Supplemental Figure 3**).

We also hypothesized that ATRX loss is a predictive biomarker for chemotherapy response given prior reports that ATRX loss is associated with greater sensitivity to DNA damaging agents (22). In the same cohort of UPS patients, responses to chemotherapy were evaluable by RECIST criteria in 35 patients, of which 16 (45.7%) had ATRX loss by IHC. In this cohort, there was no significant difference in overall response rate (ORR, 26.7% with ATRX loss, 34.2% with ATRX intact, p = 0.76) (**Supplemental Figure 4**).

Based on this clinical observation that ATRX loss is associated with worse outcomes in multiple sarcoma subtypes, we set out to better understand the mechanisms underlying this phenomenon by developing *in vitro* and *in vivo* STS models of ATRX loss.

### Loss of TP53 is required for proliferation without ATRX

To model the biology of *ATRX* loss in sarcomas, we employed primary human cell strains from the hypothesized cells of origin for each of the relevant sarcoma subtypes. The cell of origin for LMS is a smooth muscle cell or precursor mesenchymal stem cell (27), while the cell of origin for UPS is unknown though is most likely a mesenchymal stem cell (28). To model LMS, we employed a human uterine smooth muscle strain (HUtSMC), and for UPS and DDLPS we employed human mesenchymal stem cells (MSCs) and fibroblasts (BJ and HDFn). Notably, BJ, HDFn, and HUtSMC cell strains are all telomerase-negative. *ATRX* loss is associated with activation of an alternative telomere maintenance mechanism termed ALT (29–31), whereby homologous recombination (HR) machinery leads to telomere elongation in lieu of telomerase. Therefore, modeling ATRX loss with a telomerase-negative cell line may better recapitulate disease biology. CRISPR-mediated knockout of *ATRX* (**Figure 2A-B**) failed to generate monoclonal *ATRX* knockout cells (**Supplemental Figure 5**), and polyclonal cell populations were less proliferative (**Figure 2C-E**). This is consistent with previous reports that *ATRX* mutation or deletion, without co-incident tumor suppressor mutations, actually impairs cell growth in certain contexts (32). Biallelic disruption of *TP53* and *RB1* occur in 92% and 94% of LMS (12), and examination of these alterations in the sarcoma TCGA and GENIE databases showed that *ATRX* alterations frequently co-occur with *TP53* alterations (**Supplemental Figure 6**). Concordantly, the DepMap data show a negative correlation for co-dependence between *ATRX* and *TP53* (correlation -0.27), suggesting that loss of *TP53* is a critical requirement for tumor fitness with *ATRX* loss (**Table S4**). Further, negative regulators of *TP53*, including *MDM2* (0.26) and *PPM1D* (0.27), show a positive co-dependency correlation with *ATRX* (**Table S4**). To test if loss of *TP53* is required for increased proliferation of *ATRX* knockout cells (33), we depleted *TP53* using CRISPR in the aforementioned cell lines (**Figure 2F**). P21, which is a critical mediator of p53-dependent cell cycle arrest in response to DNA damage (**Supplemental Figure 7**), was downregulated in these cells. *ATRX/TP53* double knockout (dKO) monoclonal cell lines could be generated and showed increased proliferation in comparison to ATRX KO (**Figure 3E; Supplemental Figure 5**), but not when compared to *TP53* KO cells (**Figure 2G-I**).

**Figure 2.**
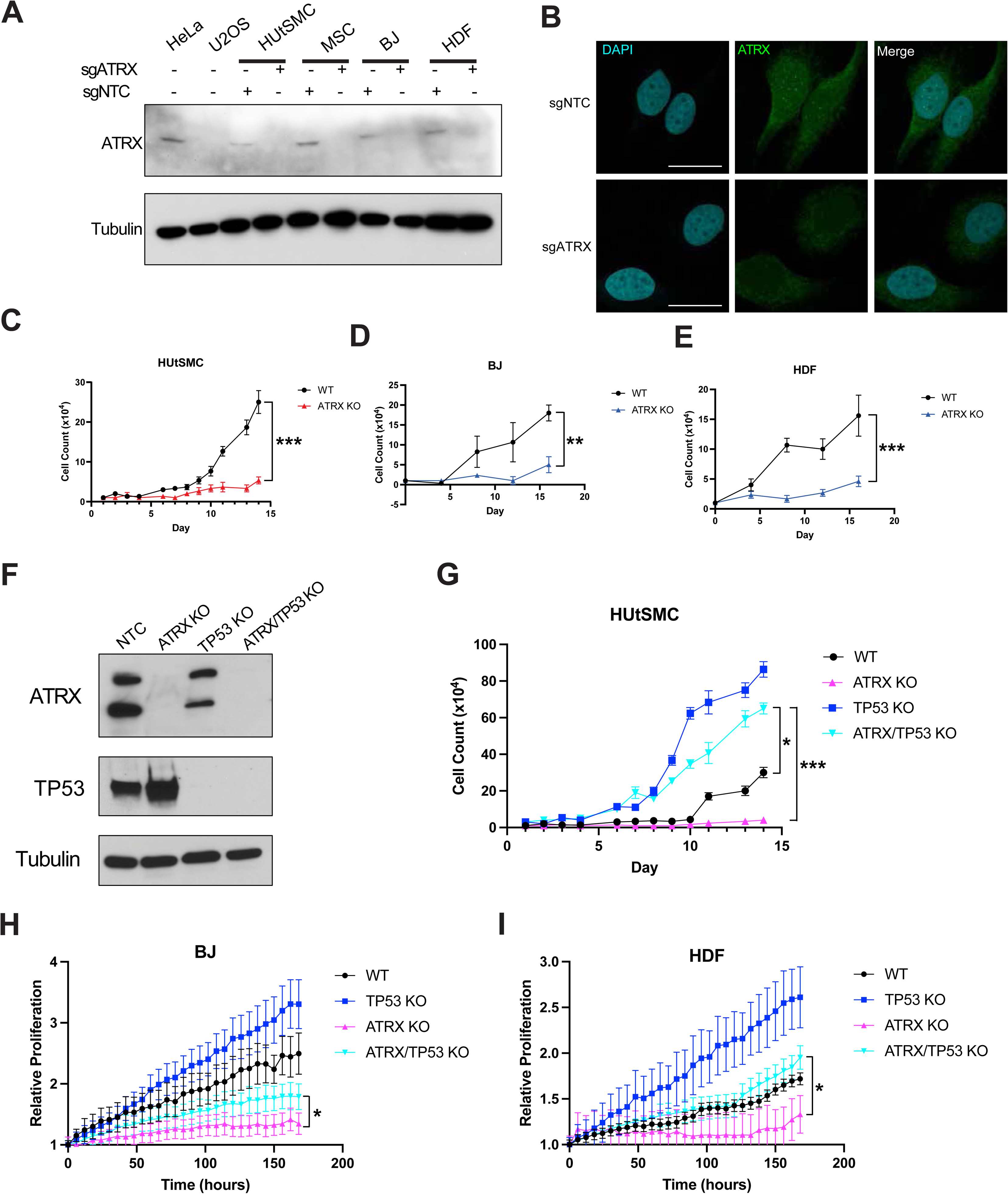
Proliferation without *ATRX* requires loss of *TP53*. (A) Western blot showing depletion of ATRX using CRISPR and sgRNA targeting ATRX or a non-targeting control sequence (NTC). (B) Immunofluorescence images of ATRX depletion. (C-E) Growth curves showing cell counts over time with ATRX knockout in HUtSMC (C), BJ (D), and HDF (E). (F) Western blots showing dual knockout of ATRX and TP53 in HUtSMC cells. (G-I) Growth curves over time showing HUtSMC cell number (G) or percent confluence for BJ (H) and HDF (I). *P value < 0.05; **P value < 0.01; ***P value < 0.001. Scale bars = 10 μm.

**Figure 3.**
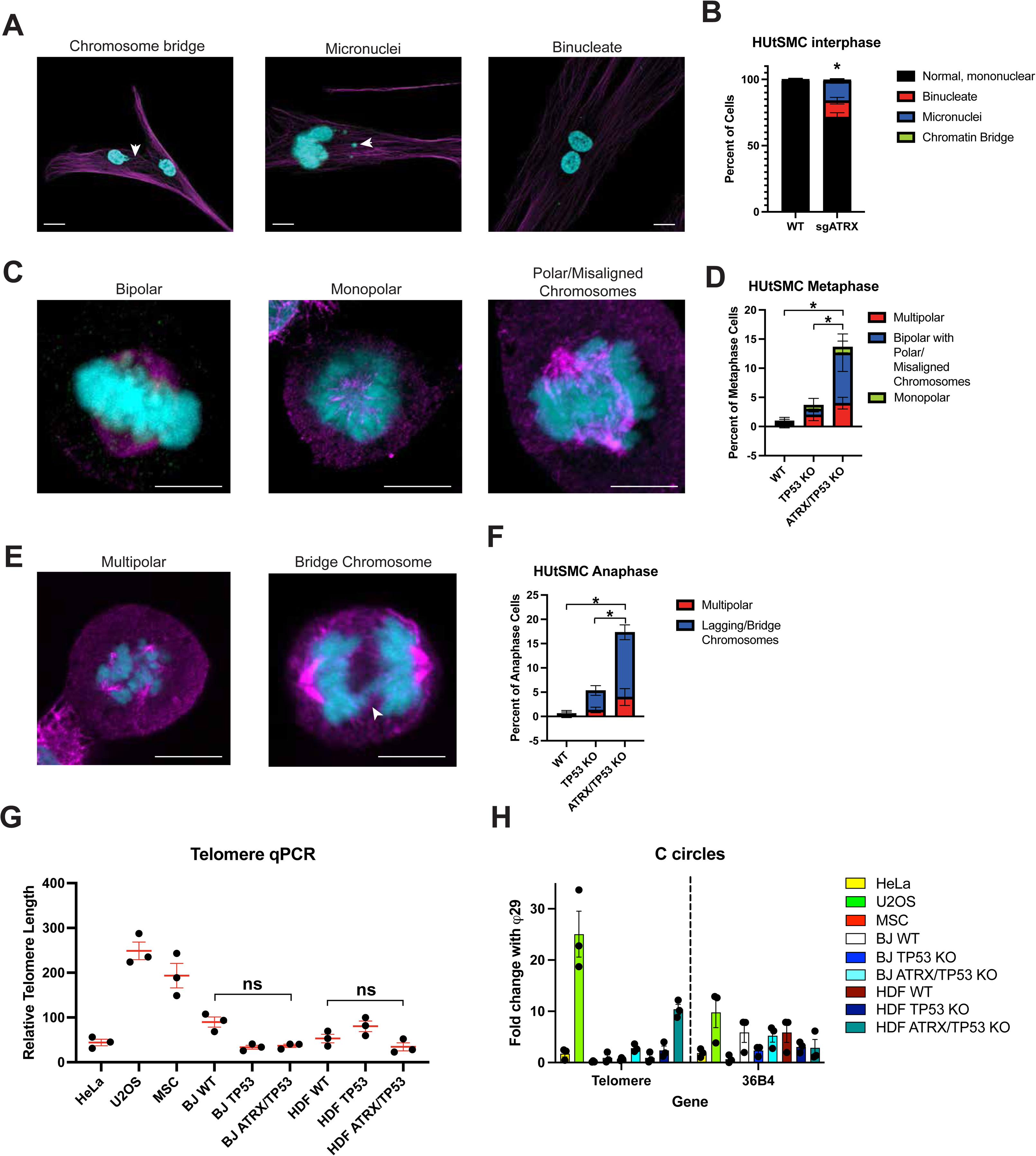
Loss of *ATRX* causes genomic instability. (A) Representative micrographs of HUtSMC *ATRX/TP53* dKO cells with a chromosome bridge, micronuclei, or multiple nuclei. (B) Quantification of these phenotypes in interphase cells. (C) Representative micrographs of HUtSMC *ATRX/TP53* dKO cells in metaphase with the indicated phenotypes. (D) Quantification of metaphase cells. (E) Representative micrographs of cells in anaphase. (F) Quantification of anaphase cells. (G) Relative telomere length, which was calculated by using the following formula: RTL = 2^−(CTtelomere−CT36B4)^, where RTL is the relative telomere length, CT_telomere_ is the cycle threshold values of telomere and CT_36B4_ is the cycle threshold values of a single-copy gene, 36B4. (H) C circle assay assessed by telomere and 36B4 quantitative PCR with and without the φ29 polymerase, which amplifies C circles. *P value < 0.05. Scale bars = 10 μm.

### ATRX loss causes genomic instability but not ALT phenotype in sarcoma

ATRX loss has previously been associated with genomic instability, and we hypothesized that one of the mechanisms by which ATRX loss contributes to worse outcomes in sarcoma is by causing genomic instability. Therefore, we assessed our cell models for evidence of genomic instability. In interphase cells, we observed an increase in binucleate cells and cells with micronuclei in *ATRX/TP53* dKO cells (**Figure 3A-B**). There were also occasional chromosome bridges (**Figure 3A**). In mitotic cells, we observed an increase in polar and misaligned chromosomes and multipolar spindles in metaphase and lagging/bridge chromosomes and multipolar spindles in anaphase (**Figure 3C-F**). By propidium iodide staining and subsequent flow cytometric analysis, we observe an increase in tetraploid (4N) cells, suggestive of cytokinesis failure (**Supplemental Figure 8**). Concordantly, in the TCGA sarcoma dataset, *ATRX* mutation was associated with significantly greater fraction of the genome altered and aneuploidy (**Supplemental Figure 8**).

Interestingly, in TCGA data, *ATRX* WT tumors are notable for increased TERT expression compared to *ATRX*-mutant tumors (**Supplemental Figure 8**), suggestive of telomerase-dependent telomere maintenance in *ATRX* WT sarcomas and ALT in *ATRX*-mutant sarcomas. As ALT+ tumors exhibit very long telomeres, we assessed telomere length in our models. HeLa cells, which express telomerase) demonstrated low relative telomere length (**Figure 3G**). Conversely, U2OS, an osteosarcoma cell line lacking ATRX expression and known to be ALT+, exhibited long telomeres (**Figure 3G**). However, we did not observe an increase in relative telomere length in our *ATRX/TP53* dKO fibroblast cell lines (**Figure 3G**) suggesting they may not have activated ALT. We also assayed for DNA C-circles, which are extrachromosomal DNA circles composed of telomeric repeats and the most specific biomarker for ALT positivity, but did not observe C circle amplification with *ATRX/TP53* KO models (**Figure 3H**). To further test this, we assessed ALT in human sarcomas using *in situ* hybridization with a telomere probe (TEL-FISH) in the aforementioned UPS TMA. In this UPS cohort (n = 72 tumors with evaluable ATRX and TEL-FISH), loss of ATRX was not significantly associated with the presence of ALT by TEL-FISH (**Supplemental Figure 9**). These findings are consistent with previously reported data that *ATRX* loss alone is insufficient for ALT (34).

### ATRX loss increases tumor growth and invasiveness

Next we asked whether *ATRX/TP53* dKO was sufficient for tumorigenesis *in vivo*. *ATRX/TP53* dKO, *TP53* KO, and WT BJ and HDFn fibroblasts were xenografted subcutaneously into the flanks of nude mice; however, this did not result in any frank tumors (data not shown). We hypothesized that additional alterations are required for tumorigenesis. Therefore, we assessed how ATRX loss affected 3 transformed sarcoma cell lines including 2 UPS cell lines (UPS186, UPS511) and one radiation-induced sarcoma (RIS620) (35, 36). All 3 of these cell lines express ATRX (**Supplemental Figure 10**) and harbor WT *ATRX* (35). We knocked out *ATRX* in these cell lines and assessed proliferation, migration, and invasion (**Figure 4A-D**). We then xenografted these cell lines to assess tumor growth *in vivo*. *ATRX* KO UPS511 tumors grew faster than *ATRX* WT UPS511 or UPS511 expressing a non-targeting control sgRNA (sgNTC) (**Figure 4E-F**). We conclude that *ATRX* loss increases tumorigenicity in transformed sarcoma cells *in vivo*.

**Figure 4.**
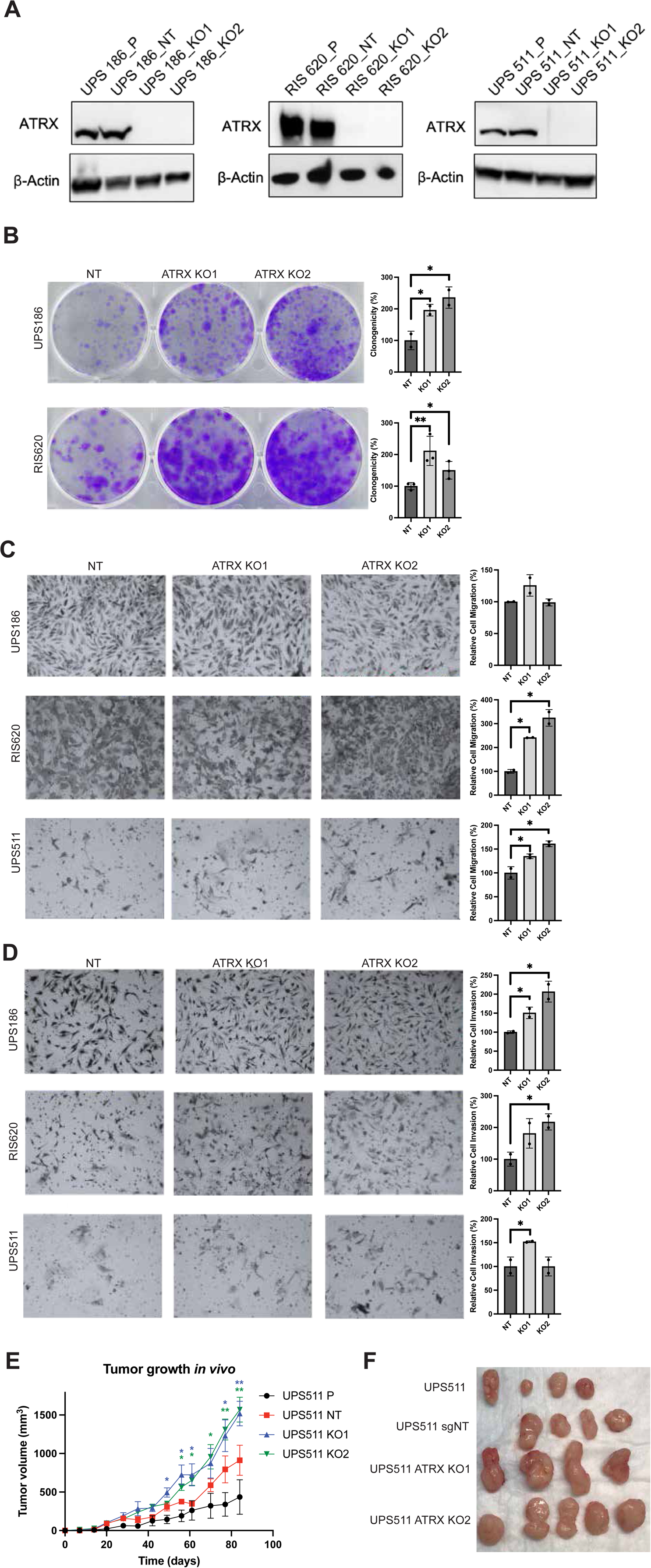
Loss of *ATRX* enhances migration, invasion, and tumorigenesis *in vivo*. (A) Western blotting for ATRX in UPS cell lines. Two different clones were used for each cell line. P indicates parental control and NT indicates expression of a non-targeting control sgRNA. (B) Representative images from colony formation assay. Quantification of crystal violet is shown in the adjacent bar graphs. (C) Representative images from migration assay. Quantification of migrating cells shown in the adjacent bar graphs. (D) Representative images from invasion assay. Quantification of invading cells shown in the adjacent bar graphs. (E) UPS cells were xenografted into immunodeficient mice, and tumor volume was measured over time. (F) Image of explanted tumors at the end of the xenograft experiment. Bars represent means ± SD, and significance is indicated as follows: *P < 0.05, **P < 0.01.

### ATRX loss causes global epigenomic reprogramming and activation of PRDM4 and NFIX transcription factors

To assess the impact of ATRX loss on the epigenome, we examined chromatin accessibility profiles through ATAC-seq in our models of ATRX loss in sarcoma precursors and in a transformed LMS cell line, Leio-012, with or without ATRX depletion. ATRX depletion led to a significant increase in the enrichment of open chromatin peaks at the TSS in Leio-012 or a *TP53* KO HUtSMC cell line, but not in the non-transformed HDF and BJ fibroblast lines (**Figure 5A**), suggesting TP53 loss may be required to permit global chromatin changes caused by loss of *ATRX*. UMAP analysis of the ATAC-seq data showed cells more closely clustered based on their cell identity or background rather than *ATRX* status (**Supplemental Figure 11**). Further, the same types of cells (e.g. BJ and HDF fibroblasts versus smooth muscle-derived HUtSMC and Leio-012 cell lines) tended to cluster together.

**Figure 5.**
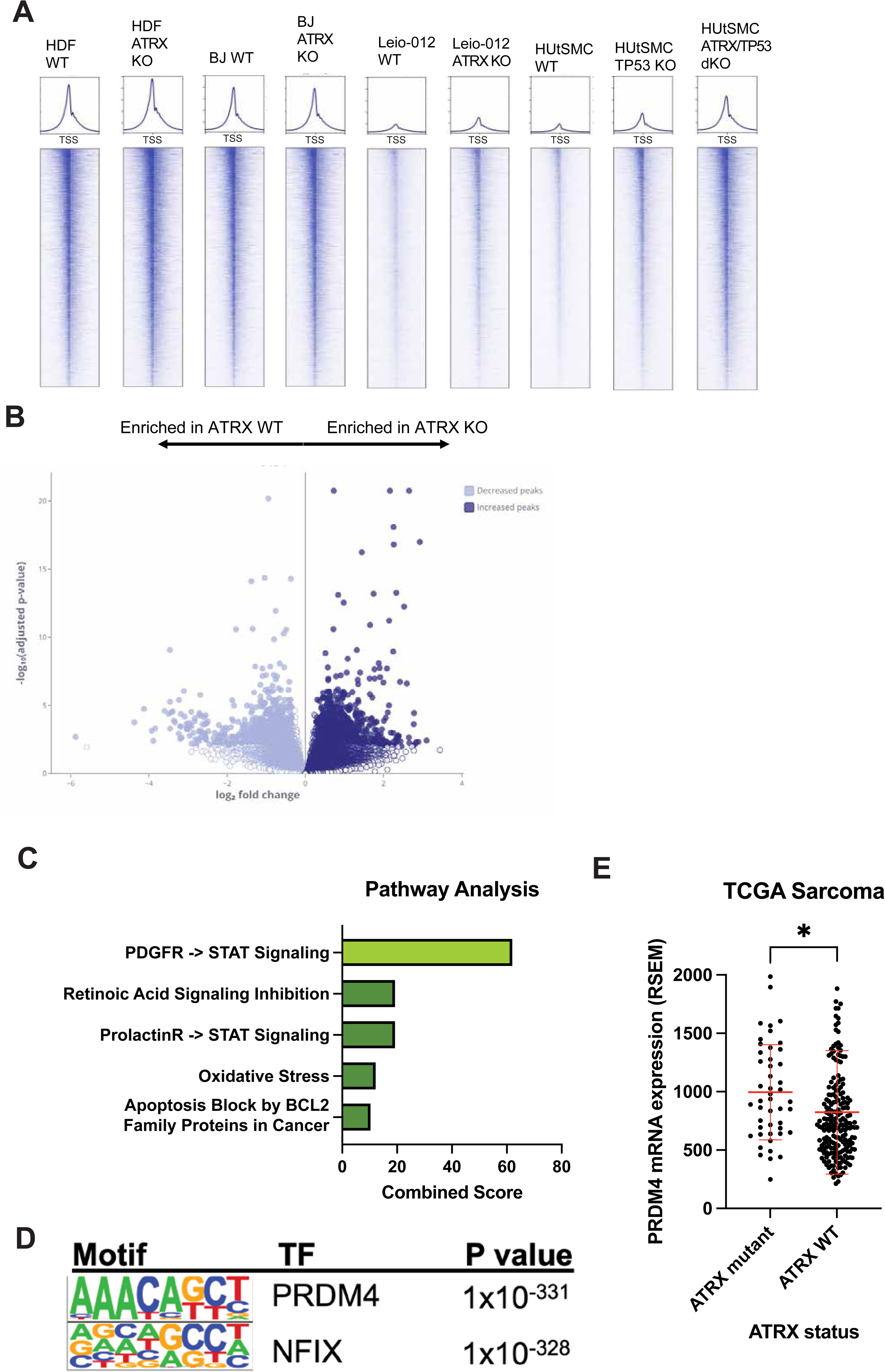
*ATRX* loss induces epigenetic rewiring. (A) Heatmap and average intensity curves of ATAC peaks centered on the TSS. (B) Volcano plot showing peaks enriched in *ATRX* KO versus WT conditions. (C) Pathway analysis of the genes corresponding to the peaks significantly enriched in *ATRX* KO. Enrichr was used for pathway analysis. (D) Homer TF motif analysis was used to identify motifs enriched in *ATRX* KO cells (versus *ATRX* WT cells). (E) *PRDM4* mRNA expression (RSEM values) in *ATRX* mutant versus WT tumors in the TCGA sarcoma dataset. Each black dot represents an individual tumor in the TCGA sarcoma dataset, and red bars represent Bars represent means ± SD, and significance is indicated as follows: *P < 0.05.

Pathway analysis of the genes corresponding to the peaks enriched in *ATRX* KO cells versus *ATRX* WT cells (**Figure 5B**) identified PDGFR-STAT signaling to be the most enriched pathway (**Figure 5C**). Some of the peaks that were most upregulated in *ATRX* KO compared to *ATRX* WT included *ZNF404* and *GALNT2* (**Supplemental Figure 11**). *ZNF404* is zinc finger protein that is mainly expressed in the brain, especially glial cells, while *GALNT2* catalyzes the initial reaction in O-linked oligosaccharide biosynthesis, the transfer of an N-acetyl-D-galactosamine residue to a serine or threonine residue on the protein receptor.

Transcription factor (TF) motif analysis using HOMER identified *PRDM4* and *NFIX* as potential TFs motifs enriched in *ATRX* KO (**Figure 5D**), and this was seen in both HUtSMC and fibroblast cell strains (**Table S5**). PRDM4 (also PFM1) is a zinc finger protein and transcription factor with a SET domain and member of the 17-member PRDM family of transcription factors, which are characterized by a positive regulatory (PR) domain and multiple zinc finger domains. PRDM4 is an HDAC-associated transcriptional repressor that affects cell cycle progression (37). In the sarcoma TCGA dataset (31), *PRDM4* amplifications and gene overexpression occur in 10.0%, while point mutations and deletions are rarely observed (**Supplemental Figure 12**). Interestingly, *ATRX* mutant sarcomas were associated with greater *PRDM4* expression (**Figure 5E**).

The other TF motif we found enriched with *ATRX* loss is *NFIX*, which is a member of the NFI family of transcription factors that also includes *NFIA*, *NFIB*, and *NFIC* (*38*). Interestingly, *ATRX* loss has been shown to upregulate a pro-astrocytic regulatory program driven by NFI genes and downregulate a pro-oligodendrocytic program driven by bHLH transcription factors (39). *NFIX* mRNA overexpression is seen in approximately 10% of sarcomas, and *NFIX* overexpression is associated with significantly improved DSS in the sarcoma TCGA (p = 0.03) though no significant difference in OS and PFS (**Supplemental Figure 12**). Compared to NFIX, other NFI members (that share same DNA binding motif) exhibited lower expression. However, in the TCGA data, there was no significant difference in NFI TF expression based on *ATRX* status (**Supplemental Figure 12**).

### *PRDM4* and *NFIX* are important for malignant properties in *ATRX*-mutant sarcoma cells

PRDM4 interacts with PRMT5, an arginine methyltransferase that mediates H4R3me2, which, among other functions, has been shown to maintain a dedifferentiated cellular state in neural stem cells (40). Further, PRDM4 was found to physically interact with YAP1 in a biochemical screen and mediates YAP1-induced cell invasion via transcription of ITGB2, an integrin subunit (41). Consistently, we observed a significant positive, though weak, correlation between *PRDM4* and *ITGB2* in the sarcoma TCGA dataset (**Supplemental Figure 13**). However, no significant association was found between *ITGB2* expression and *ATRX* status (**Supplemental Figure 13**). Given the association with PRMT5, we assessed but did not observe change in levels of H4R3me2s or H3R8me2s, suggesting no PRMT5 activation with *ATRX* loss (**Supplemental Figure 14**).

To better understand the biology of *PRDM4* and *NFIX* in the context of *ATRX*-mutant sarcomas, we employed 2 sarcoma cell lines that harbor mutant *ATRX* and do not express the protein (MPNST007 and Lipo815) and 2 matched *ATRX* WT cell lines (MPNST4970 and LPS863; **Figure 6A, Supplemental Figure 15**) (35). RNAi-mediated depletion of both *PRDM4* and *NFIX* (**Figure 6B-C**; **Supplemental Figure 16**) decreased the proliferation of all 4 cell lines (**Figure 6D-E**). However, the magnitude of proliferation decrease was more pronounced in the *ATRX* KO cell lines, Lipo815 and MPNST007 (**Figure 6F-G**). Depletion of *PRDM4* and *NFIX* also resulted in a decrease in invasion of the *ATRX* KO cell line MPNST007 (**Figure 6H-I**).

**Figure 6.**
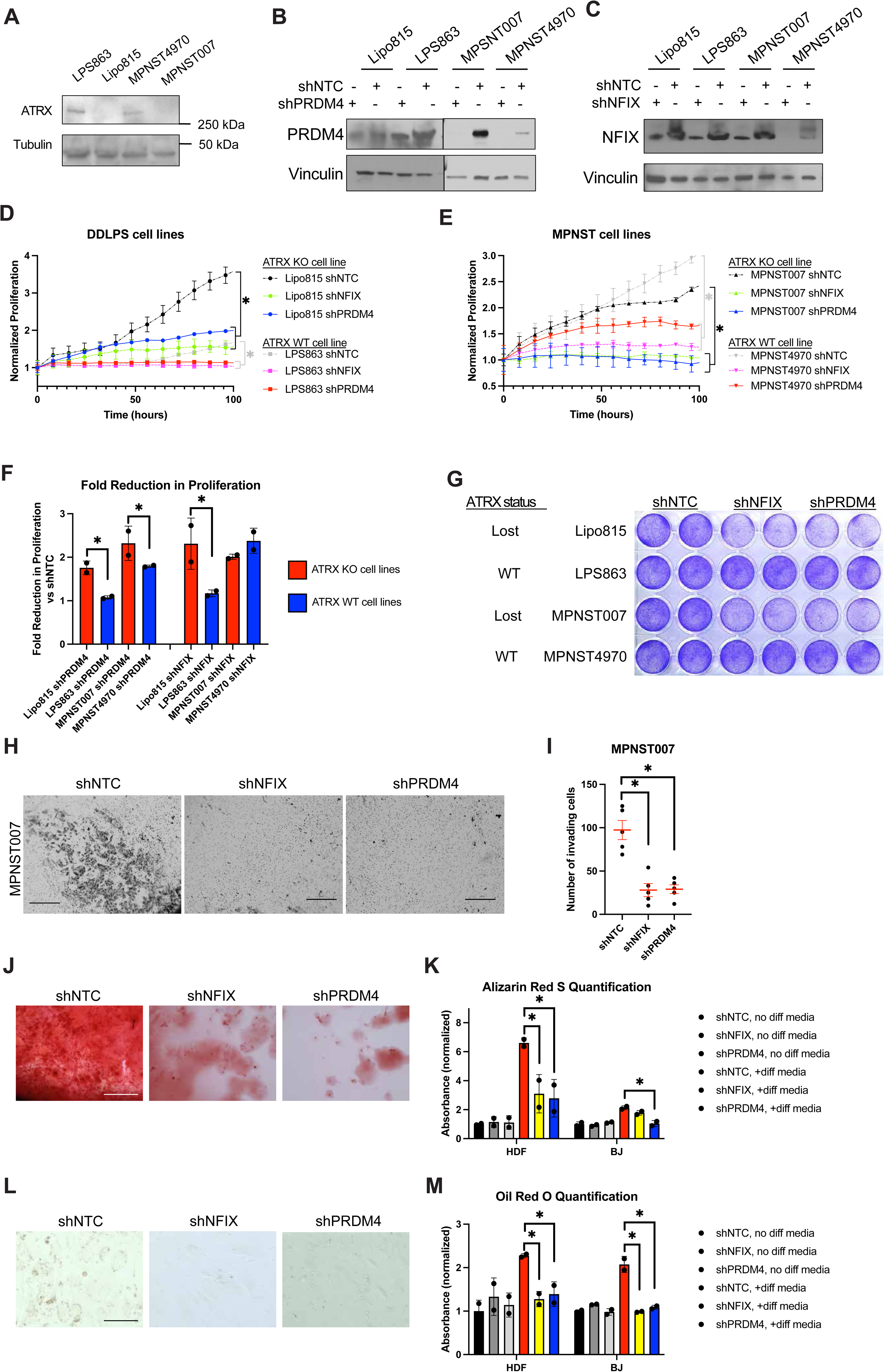
*PRDM4* depletion reduces proliferation and invasion. (A) Western blots showing ATRX expression in 2 DDLPS and 2 MPNST cell lines. (B-C) Western blots showing depletion of PRDM4 (B) and NFIX (C) using shRNA and CRISPR (sgRNAs). (D-E) Normalized proliferation determined from percent confluence. (F) Fold reduction in proliferation comparing *PRDM4* or *NFIX* depletion to non-targeting control (NTC). (G) Crystal violet assay in *ATRX* KO sarcoma cell lines (Lipo815 and MPNST007) compared to *ATRX* WT sarcoma cell lines (LPS863 and MPNST4970). (H) Representative images of crystal violet staining in invasion assay. Scale bars = 100 μm. (I) Quantification of invading cells in MPNST007 expressing the indicated shRNAs. (J) Representative micrographs of alizarin red staining. Scale bar = 10 μm. (K) Quantification of alizarin red S by eluting the dye and measuring absorbance. (L) Representative micrographs of oil red O staining. Scale bar = 10 μm. (M) Quantification of oil red O by eluting the dye and measuring absorbance. *P value < 0.05.

Next we tested if *PRDM4* and *NFIX* are important for differentiation state using BJ and HDF non-transformed fibroblasts, which are capable of differentiating into adipocytes and osteoblasts (42). Indeed, depletion of *PRDM4* and *NFIX* reduced the ability of these fibroblasts to differentiate into osteoblasts (**Figure 6J-K**) and adipocytes (**Figure 6L-M**).

### ATRX loss sensitizes to TEAD inhibition

PRDM4 binds to and activates YAP1, enhancing downstream TEAD signaling (41). Further, previous reports have shown enrichment in TEAD activity following *ATRX* KO (43). Coupled with our observations that *ATRX* KO increases *PRDM4* expression and enriches for *PRDM4* motifs, we hypothesized that increased expression of *PRDM4* drives increased YAP/TEAD activity. Therefore, we tested the hypothesis that *ATRX* loss activates YAP-TEAD signaling and confers sensitivity to TEAD inhibition. First, we validated the interaction between PRDM4 and YAP1 (and the YAP1-TEAD1-enhancer or YTE complex) by applying LPS863 shNTC and shATRX cell extracts to beads conjugated with YAP1 or YTE complex. Proteins were eluted from the beads and probed for PRDM4, which confirmed the binding of PRDM4 to both YAP1 and YAP1-TEAD1 complex (**Figure 7A**). *ATRX* depletion in the LMS cell line Leio-012 and DDLPS cell line LPS863 resulted in decreased expression of the TEAD target genes *CTGF* and *CYR61* (**Figure 7B-D**). We did not observe an increase in overall YAP1 expression with *ATRX* KO, suggesting that this effect is not dependent on YAP1 levels (**Supplemental Figure 17**).

**Figure 7.**
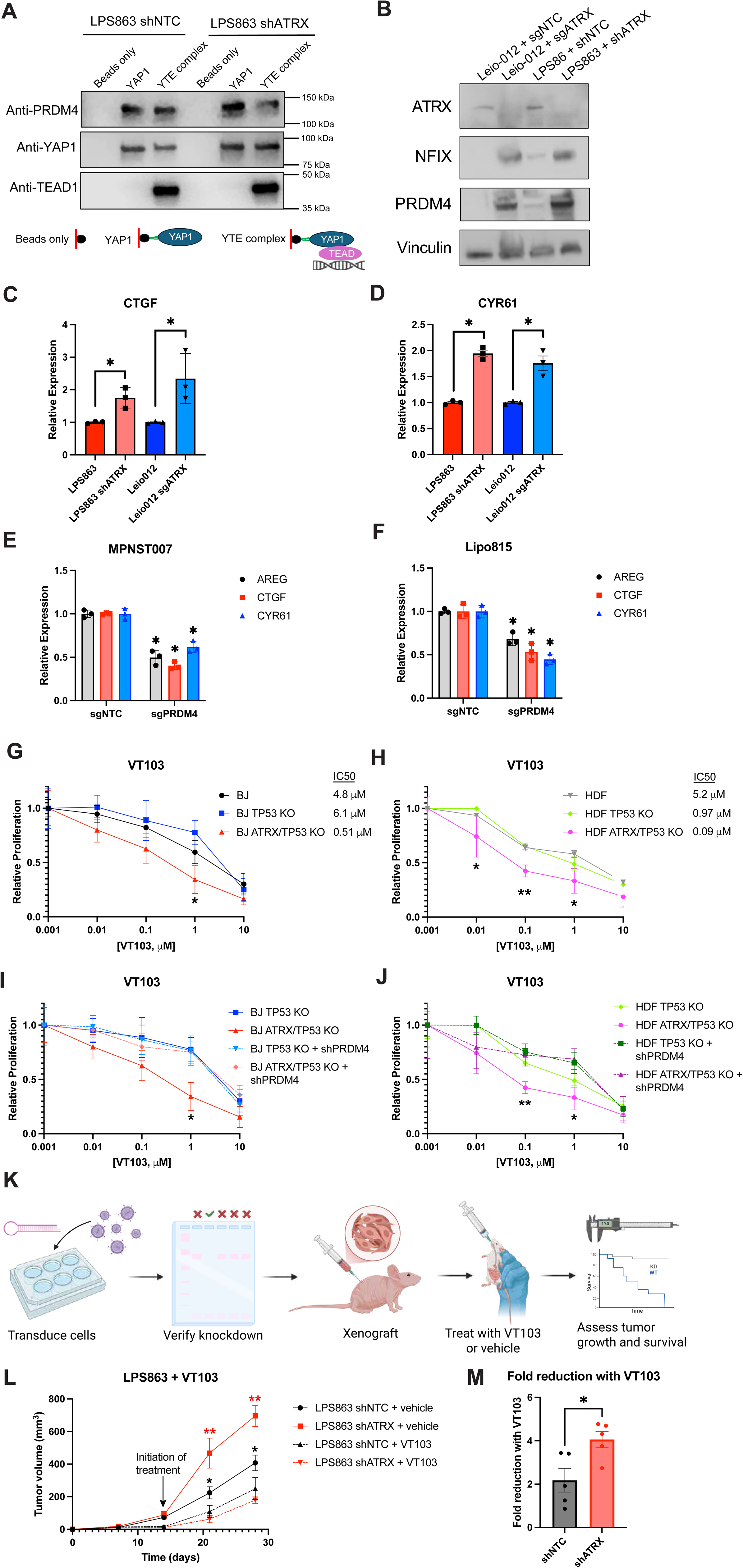
*ATRX* loss sensitizes to TEAD inhibition. (A) Western blot analysis showing that both YAP1 and the YAP1-TEAD1-enhancer (YTE) complex are able to pull down PRDM4 from LPS863. (B) Western blots showing depletion of ATRX and effect on NFIX and PRDM4 expression in Leio-012 and LPS863 cell lines. (C-D) Relative expression of *CTGF* and *CYR61* with ATRX depletion by qRT-PCR. (E-F) Expression of TEAD target genes (*AREG, CTGF*, and *CYR61*) by qRT-PCR in MPNST007 (E) and Lipo815 (F) with deletion of PRDM4. (G-H) Dose response curves with treatment with the TEAD inhibitor VT103. (I-J) Dose response curves with treatment with VT103 after depletion of *PRDM4* by shRNA. (K) Schematic of a mouse xenograft experiment testing VT103 efficacy. (L) Xenograft tumor growth curves with treatment with VT103. (M) Fold reduction comparing VT103-treated tumors to their vehicle-treated controls. Bars represent means ± SEM, and significance is indicated as follows: *P < 0.05, **P < 0.01.

Next, we tested whether depletion of PRDM4 affects YAP1-TEAD signaling using 2 sarcoma cell lines with *ATRX* loss: an MPNST cell line, MPNST007, and a DDLPS cell line, Lipo815. Depletion of *PRDM4* resulted in a decrease in the expression of TEAD target genes *CTGF* and *CYR61* (**Figure 7E-F**).

Next, we tested the sensitivity of *ATRX* KO cells to TEAD inhibition. In BJ and HDF fibroblasts, *ATRX/TP53* KO cells were more sensitive to the TEAD inhibitor VT103 compared to *TP53* KO and parental controls (**Figure 7G-H**). We next asked whether this effect depends on PRDM4. To that end, we depleted *PRDM4* from *ATRX* KO Leio-012 and *ATRX* KD LPS863 cells and tested sensitivity to TEAD inhibition. Depletion of *PRDM4* reduced the efficacy of TEAD inhibition (**Figure 7I-J**), suggesting that the sensitivity to TEAD inhibitors in the context of *ATRX* loss depends on upregulation of *PRDM4*. To test the *in vivo* efficacy of TEAD inhibition for *ATRX*-mutant sarcomas, we xenografted LPS863 cells into immunodeficient nude mice (**Figure 7K**). Unfortunately, our efforts to xenograft BJ, HDF, and Leio-012 cells into nude mice were unsuccessful. Consistent with our UPS511 xenograft model, *ATRX*-deficient LPS863 cells grew tumors at a faster rate than non-targeting control LPS863 cells (**Figure 7L**). With treatment with VT103, *ATRX*-deficient LPS863 xenograft tumors were more sensitive to VT103 and exhibited a greater fold reduction in tumor growth with VT103 (**Figure 7M**).

We conclude that *ATRX* loss causes chromatin remodeling, increases PRDM4 expression and TF activity and downstream YAP1-TEAD signaling, and confers sensitivity to TEAD inhibition.

## DISCUSSION

Our study provides several major advances in ATRX biology. First, we demonstrate that *ATRX* loss is a prognostic biomarker for poor outcomes in multiple sarcoma subtypes. Second, ATRX promotes proliferation in cooperation with p53 loss through modulation of open chromatin regions via its chromatin remodeling activity. Third, we identify reprogramming of open chromatin directed at PRDM4- and NFIX-mediated transcriptional programs. Lastly, we discover ATRX-PRDM4-YAP1 as a novel mechanistic axis that drives malignant phenotypes and is therapeutically targetable by TEAD inhibitors. This study suggests a novel precision medicine strategy to treat *ATRX* -mutant sarcomas with TEAD inhibitors.

Sarcoma treatment is largely not informed by molecular subtyping. Furthermore, there is a dearth of prognostic and predictive biomarkers. Our data support the use of ATRX expression as a prognostic biomarker for worse outcomes in multiple sarcoma subtypes. Previous data also support this notion. *ATRX* mutation/loss has been associated with worse disease-specific survival in soft tissue sarcomas in TCGA (22). In a cohort of 128 soft tissue sarcomas with multiple subtypes, reduced ATRX expression by IHC was associated with worse outcomes (44). There was also a trend toward worse OS in MSKCC cohorts of LMS and osteosarcoma though no difference in UPS and ULMS (45). *ATRX* mutation has also been associated with worse outcomes in a cohort of DDLPS (46). *ATRX* mutation has also been incorporated into a risk stratification score for LMS (47). However, ATRX IHC in a cohort of 91 LMS tumors showed that *ATRX* loss was not associated with worse survival (48). In other cancer types, *ATRX* and *DAXX* mutations have been associated with reduced survival in pancreatic neuroendocrine tumors (19, 49), while *ATRX* mutation is associated with better OS in IDH-mutant gliomas (20).

What are the mechanisms by which *ATRX* loss leads to worse outcomes? One possible explanation is increased genomic instability. There are multiple mechanisms by which *ATRX* loss leads to genomic instability. First, *ATRX* loss is associated with ALT, and telomere dysfunction has been associated with genomic instability (50, 51). Second, *ATRX* loss is associated with an increase in G quadruplex structures, which can lead to stalled replication forks.(52, 53) Third, ATRX loss is associated with impaired chromosome cohesion and congression (54). A second major explanation for worse outcomes in *ATRX*-mutant tumors is the association with a more immunosuppressive tumor microenvironment, as we show herein and as was demonstrated in ATRX KO murine gliomas with greater enrichment of immunosuppressive macrophages (39). A fourth potential mechanism is increased NFkB pathway activation and alteration of ECM (55). A fifth hypothesis is epigenetic de-repression and expression of endogenous retroviruses (ERVs), which make up 8% of the genome and have been implicated in suppression of host immunity (56). *ATRX* and *DAXX* have been shown to silence these regions of the genome (5, 57, 58).

This work has identified and implicated a novel ATRX-PRDM4-YAP1-TEAD axis in driving the growth of ATRX-mutant sarcomas. PRDM4 is zinc finger protein and TF with SET domain that is highly conserved across mammals. PRDM4 directly interacts with the YAP1 WW domains and mediates YAP1-induced cell invasion (41). In cancer, *PRDM4* expression is associated with worse outcomes in hepatocellular carcinoma and melanoma (59). Further, *PRDM4* expression is higher in metastatic prostate cancer compared to localized prostate cancer and normal prostate tissue (41). Overlap of PRDM4- and YAP-mediated genes show enrichment in mitosis, cell cycle, DNA replication, cell differentiation, and liver regeneration. Our data add to this body of literature by demonstrating that *ATRX* loss can increase *PRDM4* expression and openness of *PRDM4* motifs. Our data coupled with data from TCGA demonstrate that *ATRX* loss increases *PRDM4* expression. One possible mechanism to explain this is that *ATRX* directly binds and represses sites of *PRDM4* -mediated transcription. A second possibility is that *ATRX* loss creates abnormal transcriptional condensates at sites of *PRDM4*-mediated transcription (60). A third possibility is that there is some other indirect mechanism leading to *PRDM4*-mediated transcription. Regardless, the result of this increased *PRDM4* expression and activity is activation of YAP1-TEAD signaling, which has been heavily implicated in driving tumor growth (61, 62).

The YAP-TEAD axis promotes tumor growth through multiple mechanisms including stimulation of cell proliferation, maintaining stemness, and reprogramming metabolism, among others. Activation of this pathway may also be partially responsible for the genomic instability we observed with ATRX loss. First, YAP-TEAD promotes proliferation by increasing CDK4/6 activity, which can lead to more replication stress, DNA breaks, and genomic instability. Second, YAP1 regulates ATR/CHK1 signaling and vice versa, which can affect DNA damage (63). Lastly, YAP1 can hyperactivate the spindle assembly checkpoint and induce mitotic defects (64).

ATRX loss has been associated with ALT. Telomere maintenance is a hallmark of cancer that most often occurs by activating telomerase expression (∼90%) though can also arise by ALT (10%), a telomerase-independent mechanism. ALT-positive cells are characterized by long and heterogeneous telomeres, while ALT-negative cells generally have shorter and more homogeneous telomeres. ALT is generally more prevalent in sarcomas than carcinomas and other cancers (65–67) and may be associated with worse survival in some subtypes such as DDLPS (66). A study of 519 sarcoma samples showed significant association with loss of ATRX and presence of ALT (29). In the TCGA, a method to infer telomere length from whole exome sequencing data called TelSeq demonstrated telomere lengthening in some cases of LMS and UPS and association with *ATRX* deletion or mutation (25). In our study, we did not identify a significant correlation with ATRX loss by IHC and ALT assessed by large telomere foci on telomere FISH. Further, we did not find evidence of ALT in our *ATRX* KO fibroblast cell lines. There are several potential explanations why ALT did not correlate with ATRX loss. First, the ALT and ATRX IHC assays may not be sufficiently sensitive. Second, *ATRX* mutation does not always result in ALT in the published literature (68, 69). Third, we used cell models that were not primary human tumor-derived. In summary, our data suggest that *ATRX* loss likely facilitates but is not sufficient for ALT.

Our study has several limitations. First, IHC may not always correlate perfectly with loss-of-function, as some *ATRX* loss-of-function mutations may result in production of non-functional protein. Second, many *TP53* mutations in sarcoma are point mutations that result in expression of a mutant form that may have a dominant negative or gain-of-function effect, and point mutations have been shown to be more efficient in driving sarcoma development in mouse models (70). However, we utilized CRISPR to knockout *TP53*, which may not perfectly recapitulate *TP53* biology in sarcoma.

In summary, we identify ATRX loss as a prognostic biomarker of worse outcomes in multiple sarcoma subtypes, implicate a novel ATRX-PRDM4-YAP1-TEAD axis driving the pathogenesis of *ATRX*-mutant sarcomas, and identify a possible precision medicine strategy to target *ATRX*-mutant sarcomas.

## METHODS

### Sarcoma tissue microarrays (TMAs) and immunohistochemistry

Data from these sarcoma microarrays have been previously published (71, 72). ULMS, STLMS, UPS, and DDLPS TMAs were stained for ATRX (polyclonal, Sigma-Aldrich, 1:400) using a Leica BOND III auto stainer (Leica Biosystems, Buffalo Grove IL). The Bond Polmer Detection kit with DAB (Leica Biosystems) was used for primary antibody detection. ATRX was considered intact if the tumor had nuclear staining of any extent and of any intensity. Complete absence of nuclear staining in tumor cells was defined as complete loss of ATRX expression (with retained staining in normal cells). As a positive control in the ULMS TMA, 18 non-LMS samples including 9 uterine leiomyomas, 5 normal myometrium, and 4 esophageal leiomyomas stained positive for ATRX.

For the UPS cohort, each patient’s imaging was assessed for response using RECIST 1.1 criteria. Best responses were compared between ATRX loss and ATRX WT groups for percent change as well as response rates (CR/PR/PD/SD).

### Cell Culture

Human primary bone marrow-derived mesenchymal stem cells (MSCs) were isolated from healthy donors and provided through an IRB-approved partnership with the Texas A&M Institute for Regenerative Medicine. MSCs were selected based on positivity for CD90, CD105, CD73 127 and negativity for CD34, CD19, CD11b, CD45, CD79a, CD14, HLA-DR, HLA-DQ, and HLA-DP. These cells were cultured in minimum essential media (HyClone MEM with 2% L-glutamine) supplemented with 10% FBS and 1% non-essential amino acids. Primary human uterine smooth muscle strain HUtSMC (PCS-460-011) and primary human fibroblast strains BJ (CRL-2522) and HDFn (PCS-201-010) were obtained from ATCC. All cells were found to be negative for Mycoplasma (abm) and were tested routinely. LPS224 and LPS863 (both patient-derived dedifferentiated liposarcoma) cell lines were obtained from the MDACC Cytogenetics and Cell Authentication Core (73). MPNST007, MPNST4970, MPNST3813E, MPNST642, S462, MPNST724, and T265 were derived from human malignant peripheral nerve sheath tumors (35). ST88-14 cells were provided by Dr. Jonathan Fletcher (Brigham and Women’s Hospital, Boston, MA), and S462 cells were provided by Dr. Lan Kluwe (University Medical Center Hamburg-Eppendorf). Leio-012 cell were derived as described (74). HUtSMC were grown with DMEM/F-12 (Corning 10-090-CV) supplemented with 10% FBS (Gibco) and 1% penicillin/streptomycin (HyClone, SV30010). All other cell lines were grown in DMEM (HyClone, SH30243.01) with 10% FBS and 1% penicillin/streptomycin. STR analysis was performed on all cell lines to confirm identity.

Cellometer Mini (Nexcelom) was used for counting cells to ensure equal cell counts for experiments.

For proliferation and growth curves, an Incucyte was used (Sartorius). For invasion assays, 5x10^4^ cells were seeded in serum-free media onto BioCoat Matrigel Invasion Chambers (Corning, 8 μm pore). Complete media (10% FBS) was added into well below the chamber. After 24 hour incubation, a cotton swab was used to remove cells on top of the membrane, and the cells on the lower surface of the membrane were stained with crystal violet and quantified by microscopy with manual counting.

Lentiviruses were generated by transfecting a T25 of ∼70% confluent HEK-293T cells with 2 μg pLKO.1 vector, 1 µg pVSV-G, 1 µg psPAX2, and Lipofectamine 3000 (Thermo Fisher). Fresh medium was applied at 24 hours post-transfection, and virus was harvested 24 hours later, clarified by centrifugation at 1000g x 5 minutes and filtration through a 0.45 μm filter to remove cell debris, and diluted 1:4 with complete medium containing 10 μg/mL polybrene. Target cells were transduced at ∼50% confluence for 24 hours, then given fresh media for 24 hours, then 1 µg/mL puromycin was added for 3-5 days.

Chemicals used in this study include: puromycin (InvivoGen), polybrene (InvivoGen), propidium iodide (Thermo Fisher), and zeocin (Thermo Fisher).

### Cell Cycle Analysis

To determine cell cycle and DNA content, cells were harvested, washed in PBS, fixed in 70% ethanol until all samples were collected for analysis, then stained overnight on ice with propidium iodide (50 μg/mL PI and 100 μg/mL RNase A in PBS) followed by flow cytometric analysis with a LSRFortessa cytometer (BD Biosciences) at the MDACC Flow Cytometry and Cellular Imaging Core. The percentages of cells in G_1_-, S-, and G_2_–M phases were determined using the cell-cycle analysis program Modfit LT (Verity Software House).

### Adipogenic and Osteogenic Differentiation

Fibroblasts and MSCs were plated in 24-well plates and grown to 80–90% confluency. Adipogenic or osteogenic differentiation media (Gibco StemPro adipogenesis and osteogenesis differentiation kits) was added and changed every 3–4 days for a total of 21 days. Adipocyte lipid droplets were detected by oil red O staining (Sigma-Aldrich). Osteoblast calcification was detected by alizarin red S staining (Sigma-Aldrich). After fixation with 4% PFA, adipocyte lipid droplets were detected by oil red O staining (Sigma-Aldrich, 0.35g in 100 mL isopropanol, filter 0.2 μm, combine 6mL of this stock with 4mL water). Oil red O was quantified by eluting with 100% isopropanol and assessing absorbance at 500nm. Osteoblast calcification was detected by alizarin red S staining (Sigma-Aldrich, 1g in 50mL water, adjusted pH to 4.2, filter 0.2 μm) after fixation with 100% ethanol. Alizarin red S was quantified by eluting with mixture of 20% methanol and 10% acetic acid in water and assessing absorbance at 450nm.

### Molecular Biology

Human shRNAs (Horizon GIPZ Lentiviral shRNA) and cDNAs (Horizon Precision LentiORF collection) for the following genes were obtained from the MDACC Functional Genomics Core: PRDM4 (clone ID V3LHS_307343, mature antisense TCTTCTTTTGACATACGCT), NFIX (clone ID V3LHS_343918, mature antisense TTGATTGTGACTCCAATGT), and non-targeting control (mature antisense ATCTCGCTTGGGCGAGAGTAAG). Plasmids were used to generate lentivirus, as described above.

We obtained 3 *ATRX* CRISPR all-in-one lentivector sets (abm,12837111) with gRNAs targeting: 106: CCACGACTTGCAATGAATCA; 1422: GCATCAGAATGTTCCAACAG 1507: CAATATGAACCTGCCAACAC. To knockout *TP53*, we used pLentiCRISPR_zeo_sgTP53 (Addgene 183180).

### Western blotting

Cells were lysed in RIPA buffer (Boston Bioproducts, BP-115) with protease and phosphatase inhibitors (Halt, Thermo, 1861282). Proteins were separated by SDS-PAGE (Biorad pre-cast gels), transferred to PVDF membrane (Biorad Trans-Bot turbo 0.2μm PVDF) using Biorad Trans-Blot Turbo Transfer System, blocked with Biorad EveryBlot Blocking Buffer for 5-10 minutes, and probed overnight in 4°C with primary antibodies diluted in EveryBlot Blocking Buffer. Horseradish peroxidase-conjugated rabbit and mouse secondary antibodies (CST 7074 and 7076) were applied for 2-3 hours at room temperature. Primary antibodies used include: ATRX (Santa Cruz, D-5, sc-55584), TP53 (Santa Cruz, DO-1), p21 (CDKN1A, Cell Signaling, 2947), p21 (Abcam, ab188224), PRDM4 (Abcam, ab126939), NFIX antibody (Thermo, MA5-946983), YAP (Cell Signaling, D8H1X, 14074), alpha tubulin (Santa Cruz sc-32293), beta actin (Sigma, A2228), vinculin (Santa Cruz sc-59803), H3R8me2s (Diagenode, C15410287), H4R3me2s (Active Motif, 61988), and SDMA (gift from Mark Bedford, PhD). We also tested the following ATRX antibodies but did not have success by western blotting and immunofluorescence: PCRP-MIS18BP1-1B10 (DSHB) and PCRP-ATRX-1E4 (DSHB).

### Quantitative reverse transcription polymerase chain reaction (qRT-PCR)

For gene expression analysis by qRT-PCR, RNA was isolated from cells using Qiagen RNeasy kit. Reverse transcription was performed using SuperScript VILO (Thermo Fisher). Samples were amplified on a QuantStudio 6 (Thermo Fisher). The average C_T_ value of three housekeeping genes (*RRN18S, GAPDH,* and *ACTB*) was subtracted from each experimental C_T_, then 2−ΔCT values were normalized to controls (non-targeting shRNA control) and compared. Additionally, the ΔΔC_T_ method was employed to calculate the fold change in gene expression. Primer sequences are provided in **Table S6**.

### Immunofluorescence (IF)

Cells were plated onto glass coverslips (MatTek, PCS-1.5-18) in 12-well plates and fixed with 4% paraformaldehyde in PBS. Fixed cells were then blocked for 30 min in 3% bovine serum albumin (BSA) and 0.1% triton X-100 in PBS (PBSTx + BSA). Primary antibodies were incubated in PBSTx + BSA for 1 h at room temperature and washed three times in PBSTx, followed by secondary antibody incubation in PBSTx + BSA for 30 min at room temperature and two washes with PBSTx. Cells were counterstained with DAPI and mounted on glass slides with Prolong Diamond antifade medium (Invitrogen). Image acquisition was performed using a Keyence BZ-X microscope and a Zeiss confocal LSM880 with Airyscan.

Antibodies used for IF include: ATRX (Santa Cruz, D-5, sc-55584), PRDM4 (Abcam, ab156867), NFIX (Thermo, MA5-946983), YAP (Cell Signaling, D8H1X, 14074), alpha tubulin (DSHB Hybridoma Product 12G10, deposited to the DSHB by Frankel, J. / Nelsen, E.M.), CREST (Calbiotech, HCT-0100), pericentrin (Abcam, ab4448). Alexa fluor-conjugated secondary antibodies were used (Invitrogen, 1:350).

### Invasion assay

Invasion was assessed using Corning BioCoat Matrigel Invasion Chambers with 8.0um pore polyester (PET) membrane in 24-well plates. 1x10^5^ cells in 100 μL serum-free media were added to the top chamber. Media with 10% FBS was placed in the well below each chamber. After 24 hour incubation at 37°C, cells on the upper surface were removed by wiping with cotton swab, and the membrane was stained with crystal violet. Images were captured using a Keyence BZ-X microscope, and invading cells were manually quantified.

### ALT Assay

Determination of ALT was performed as previously prescribed.(75) Briefly, this involves a 2-step process. In the first step, 30ng of genomic DNA is subjected to rolling circle amplification by φ29 polymerase (NEB, M0269). This is followed by telomere qRT-PCR. Non-amplified gDNA is used as a control. Fold change comparing amplified to non-amplified samples is calculated for telomeres and a single copy gene (36B4). Primer sequences are shown in **Table S6**. U2OS was used as a positive control, and HeLa was used as a negative control based on prior work.(76)

### Xenografting

All animal studies were approved by the MDACC Institutional Animal Care and Use Committee (IACUC). LPS863 cells (10 x 10^6^) were injected into the flanks of nude mice (Jax, NU/J). Calipers were used to measure tumor size, and tumor volume was calculated according to the formula (a × b^2^)/2, where “a” was the longest diameter and “b” was the shortest diameter of the tumor. Treatment was initiated when tumors reached approximately 200 mm^3^. Tumors were harvested once tumor volume reached 1,500 mm^3^ or at the end point of treatment. Mice were treated with VT103 (MedChem Express, HY-134955), which was first dissolved to a stock concentration of 2.5 mg/mL using DMSO and then given at a dose of 5mg/kg in corn oil prior to administration to mice by oral gavage (2 days on, 1 day off, repeated).

### ATAC-seq

ATAC was performed as previously described (77) using Illumina Tagment DNA Enzyme and Buffer kit (20034197). 50,000 fresh cells were harvested and incubated with cold lysis buffer (10 mM Tris-HCl pH 7.4, 10 mM NaCl, 3 mM MgCL2, and 0.1% IGEPAL CA-630). Cells were centrifuged to isolate nuclei and then subjected to a transposition reaction at 37° C for 30 minutes. Transposed DNA was purified using Qiagen MinElute Purification Kit and PCR amplified with 10 cycles. PCR product concentration and quality were assessed using a high sensitivity D1000 tape station (Agilent). Sequencing was performed on an Illumina NextSeq500 at the MDACC ATGC. Analysis and figures from ATAC-seq data were generated using Pluto (https://pluto.bio).

### YAP/TEAD binding assay

Full-length YAP1 was recombinantly expressed in *E. coli* and purified. The YAP-TEAD (YTE) complex was assembled by mixing YAP1, full-length TEAD1 (CusaBio, CSB-YP023363HUf0), and a TEAD-specific enhancer DNA fragment (1 µM : 1 µM : 150 nM), followed by 1 h incubation on ice. YAP1 carried a C-terminal FLAG tag for capture on Anti-FLAG M2 magnetic beads (Sigma; M8823). Samples prepared included: (i) beads only, (ii) YAP1 bound to beads, and (iii) YTE complex bound to beads. Beads were washed five times with YAP1 buffer (25 mM Tris-HCl, 300 mM NaCl, 5% glycerol, 5 mM β-mercaptoethanol, pH 8.0). LPS863 shNTC and LPS863 shATRX cells were lysed in YAP1 buffer containing phosphatase inhibitors (Halt, Thermo, 1861282) and 1% NP-40 detergent, incubated on ice for 1 h, and centrifuged at 21,000 × g for 15 min at 4 °C. Supernatants were applied to the beads, YAP1-beads, and YTE complex–beads. Beads were washed thoroughly with YAP1 buffer until A280 of the flowthrough was negligible, mixed with SDS-PAGE loading dye (5% β-mercaptoethanol), and heated at 95 °C for 5 min. Proteins bound to beads were separated by SDS-PAGE (Bio-Rad pre-cast gels) and transferred to PVDF membranes (Bio-Rad Trans-Blot Turbo, 0.2 µm PVDF). Membranes were blocked with Bio-Rad EveryBlot Blocking Buffer (5–10 min), incubated overnight at 4 °C with primary antibodies, and probed with HRP-conjugated rabbit secondary antibodies (CST, #7074) for 1 h at room temperature. Primary antibodies included PRDM4 (Abcam, ab126939), YAP1 (Cell Signaling, D8H1X, #14074), and TEAD1 (Cell Signaling, D9X2L, #12292).

### Clinical correlates

The AACR GENIE database (version 13.1) was evaluated for soft tissue and bone sarcomas cases.(23, 24) All AACR GENIE project data have been de-identified using the HIPAA Safe Harbor Method. Analyses were retrospective and were performed in accordance with the AACR GENIE Human Subjects Protection and Privacy policy. Institutional Review Board (IRB) details are provided in the AACR GENIE Data Guide. Data were accessed using cBioPortal (78).

For determining the most commonly mutated genes, we considered only those genes for which at least 10% of the samples within the category had been profiled. OncoKB (oncokb.org) was utilized to annotate the type and function of gene alterations and to determine actionability of each mutational event (79). Two-tailed *t* tests and one-way ANOVA were used for comparisons. For multiple comparisons, Tukey’s correction was made. P-values < 0.05 were considered significant for all tests, and designations are made in the figures for statistical significance. GraphPad Prism (version 9) was utilized for descriptive statistics.

Overall survival (OS) was calculated from the date of histologic diagnosis (either pre-treatment biopsy or surgical pathology) to death or the latest follow-up. For patients with primary/localized disease, recurrence-free survival (RFS) was calculated from the date of surgery to the date of recurrence or the latest follow-up. For patients with metastatic disease, progression-free survival (PFS) was calculated from the date of diagnosis to date or progression, death, or latest follow-up. Univariate analysis of the association between survival and *ATRX* mutation status was performed using the Kaplan-Meier method and the log-rank test in GraphPad Prism. Kaplan–Meier curves were used to compare survival among different groups. Cox proportional hazards modeling using R (version 4.2.2) packages survival and survminer was used to assess the impact of multiple variables on survival. The model included the following variables: ATRX status, patient age, stage, grade, and primary tumor size.

### Statistics

Statistical analyses were performed using GraphPad Prism (version 10.0.3 or higher, RRID:SCR_002798) and R (version 4.5.0 or higher). Survival was assessed using the Kaplan Meier method. Cox proportional hazards modeling was performed using survival package in R.

For *in vitro* experiments, the data from 3 biological replicates are presented as mean ± SD, and SD represents the deviation between the biological replicates. Graph Pad Prism and R were used for statistical analyses.

Graphics for schematics in the figures and graphical abstract were created using Biorender.com.

### Study Approval

This study was approved by The University of Texas MD Anderson Cancer Center (MDACC) Institutional Review Board (protocols DR09-0245 and LAB-04-0890) and was conducted in accordance with the U.S. Common Rule. Electronic medical records were reviewed for: demographics, family history, germline testing, and characteristics of the sarcoma (e.g. histologic subtype, location, size, mitotic rate, somatic sequencing results), and sarcoma treatment history.

### Data Availability

The data leading to the reported findings in this paper are deposited in GEO. Other data are available upon request from the corresponding author.

## Acknowledgements

The authors acknowledge support from the MD Anderson Cancer Center Support Grant (P30 CA016672), the QuadW Foundation, the National Leiomyosarcoma Foundation, NIH grants T32 CA009666, 5K12CA088084, and P50CA272170, and a Conquer Cancer - Endowed Young Investigator Award in Honor of Grant R. and Victoria A. Merryman. The content is solely the responsibility of the authors and does not necessarily represent the official views of the National Institutes of Health. Any opinions, findings, and conclusions expressed in this material are those of the author(s) and do not necessarily reflect those of the American Society of Clinical Oncology® or Conquer Cancer®. The results shown here are in part based on data that were generated by the TCGA Research Network: http://cancergenome.nih.gov/. We acknowledge the TCGA Research Network, including the specimen donors and research groups, for their contributions. The authors would like to acknowledge the American Association for Cancer Research and its financial and material support in the development of the AACR Project GENIE registry, as well as members of the consortium for their commitment to data sharing. Interpretations are the responsibility of the study authors. SML and ADB were supported by the Jay Vernon Jackson Liposarcoma Fund Foundation.

