## Supplemental Figures for "ATRX Loss Predicts Poor Outcomes and Reveals a Therapeutic Vulnerability to TEAD Inhibition in Soft Tissue Sarcomas"

**Supplemental Figure 1. Most commonly altered genes in sarcoma in GENIE database.**

Bar graphs show the most common genes with mutations (A), copy number variations (B), and structural variants (C) in sarcoma samples in the GENIE database. Percent of profiled samples are shown. A gene must have been profiled in at least 10% of samples within the group (e.g. ST URS) to be included on the graph. Only those genes that were altered in at least > 5% of profiled sarcomas within the group are shown. In (B), deletions are denoted with blue bars, while amplifications are denoted with red bars.

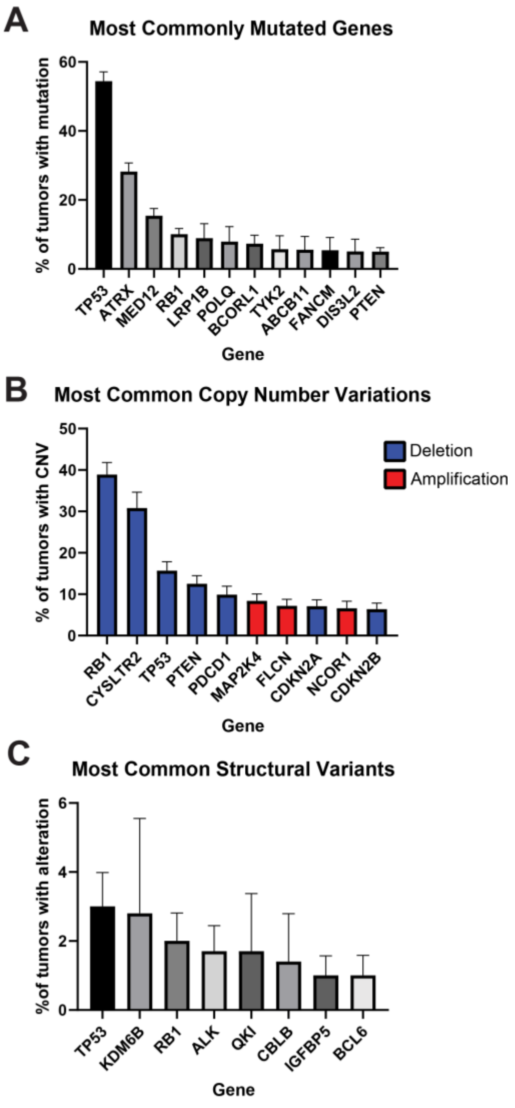

**Supplemental Figure 2. Association of ATRX alteration with survival in TCGA sarcoma dataset.**

(A-D) Overall survival (A), progression-free survival (B), disease-specific survival (C), and disease-free survival (D) in the entire sarcoma TCGA cohort based on *ATRX* alteration (red lines for mutant, black for wild type). (E-F) Overall survival (E) and progression-free survival (F) for just the patients with LMS in the TCGA sarcoma study. (G-H) Overall survival (G) and progression-free survival (H) for just the patients with UPS in the TCGA sarcoma study. (I-J) Overall survival (I) and progression-free survival (J) for just the patients with DDLPS in the TCGA sarcoma study.

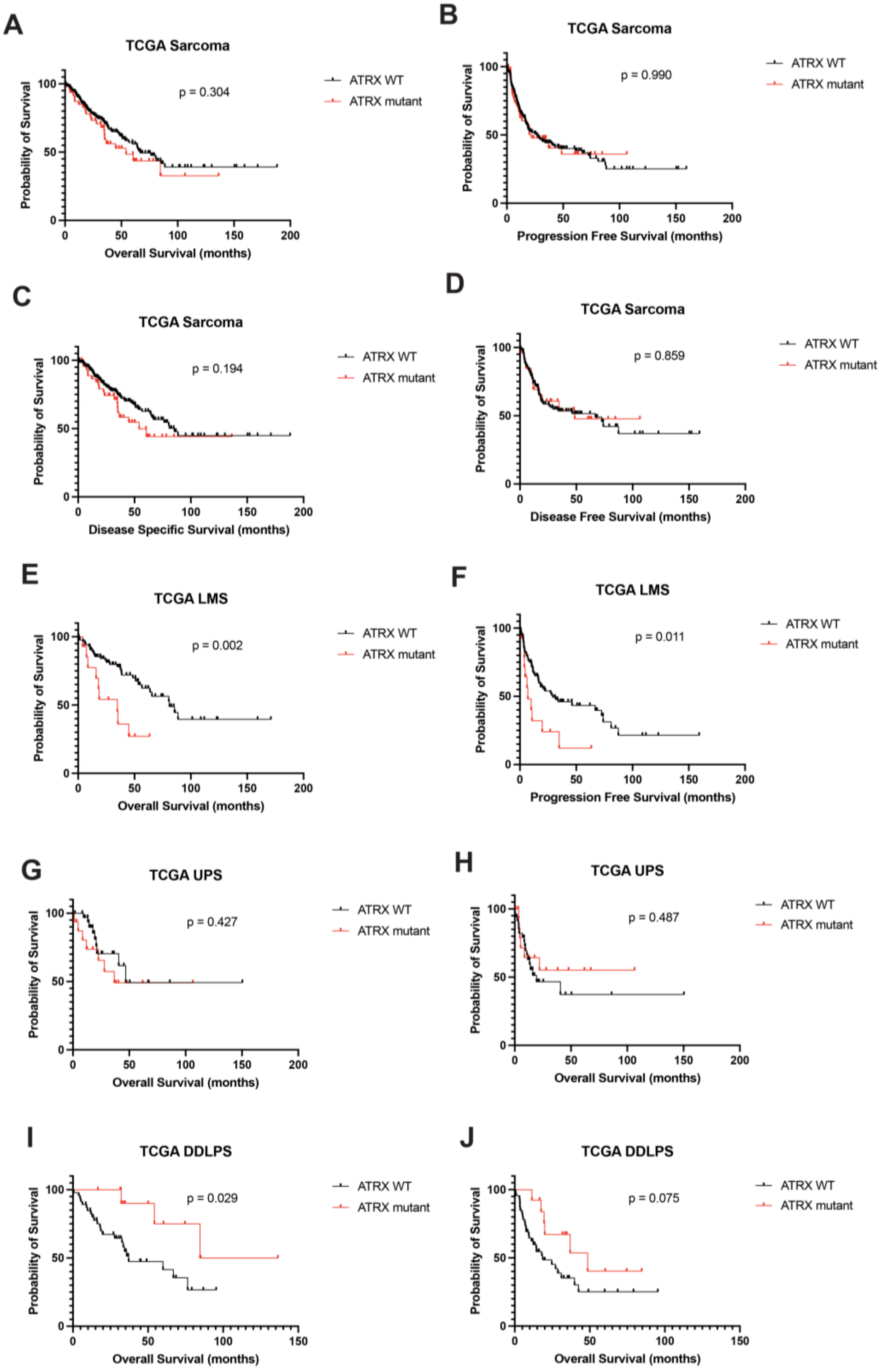
A
TCGA Sarcoma
p = 0.304
ATRX WT
ATRX mutant
Probability of Survival
Overall Survival (months)
B
TCGA Sarcoma
p = 0.990
ATRX WT
ATRX mutant
Probability of Survival
Progression Free Survival (months)
C
TCGA Sarcoma
p = 0.194
ATRX WT
ATRX mutant
Probability of Survival
Disease Specific Survival (months)
D
TCGA Sarcoma
p = 0.859
ATRX WT
ATRX mutant
Probability of Survival
Disease Free Survival (months)
E
TCGA LMS
p = 0.002
ATRX WT
ATRX mutant
Probability of Survival
Overall Survival (months)
F
TCGA LMS
p = 0.011
ATRX WT
ATRX mutant
Probability of Survival
Progression Free Survival (months)
G
TCGA UPS
p = 0.427
ATRX WT
ATRX mutant
Probability of Survival
Overall Survival (months)
H
TCGA UPS
p = 0.487
ATRX WT
ATRX mutant
Probability of Survival
Overall Survival (months)
I
TCGA DDLPS
p = 0.029
ATRX WT
ATRX mutant
Probability of Survival
Overall Survival (months)
J
TCGA DDLPS
p = 0.075
ATRX WT
ATRX mutant
Probability of Survival
Overall Survival (months)

**Supplemental Figure 3. ATRX loss is associated with worse outcomes in multiple sarcoma subtypes.**

(A) Recurrence-free survival for primary localized STLMS. (B) Overall survival for recurrent/metastatic STLMS. (C) Progression-free survival for recurrent/metastatic STLMS. (D) Recurrence-free survival for primary localized UPS. (E) Overall survival for recurrent/metastatic UPS. (F) Progression-free survival for recurrent/metastatic UPS. (G) Overall survival for DDLPS. (H) Progression-free survival for DDLPS. (I) Percent of tumor cells with TP53 positive nuclei. (J) TP53 intensity by IHC.

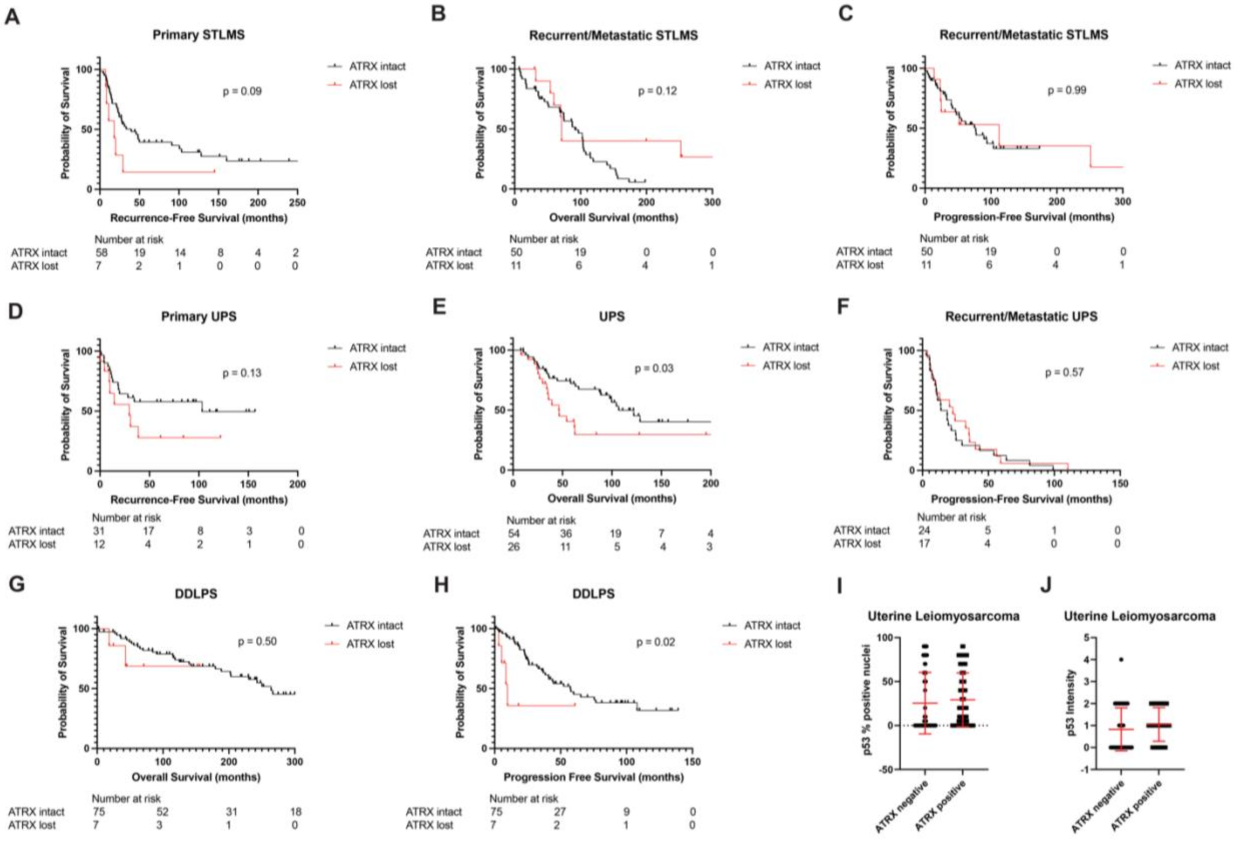

**Supplemental Figure S4. Assessment of ATRX as a predictive biomarker in human UPS.**

(A) Stacked bar plot showing proportion of patients with complete response (CR), partial response (PR), progressive disease (PD), or stable disease (SD) to chemotherapy by RECIST criteria based on ATRX status by IHC. (B) Waterfall plot showing best response to chemotherapy based on ATRX status by IHC.

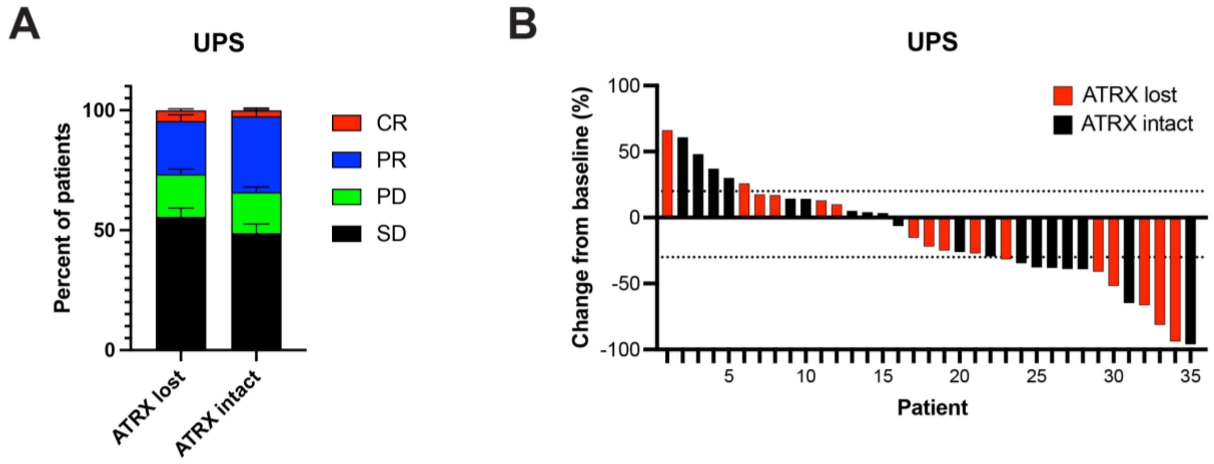

**Supplemental Figure S5. Generation of ATRX/TP53 double knockout cells.**

(A) *ATRX* is located at chromosome Xq21.1, spans 281kb, has 40 exons with ~10 different protein-coding transcripts, though the *ATRX* canonical sequence (P46100-1 in Uniprot) has 2492 amino acids, 35 exons. (B) Schematic of CRISPR and assessment of cloning efficiency. CRISPR was used to knockout *ATRX*. Several gRNAs were tested by surveyor assay and western blotting. Pools of cells with successful knockout of *ATRX* were subcloned by limiting dilution. Colonies were expanded and screened by immunofluorescent staining for *ATRX*. Made with Biorender. (C) Cloning efficiency for *ATRX* knockouts. Cells were plated by limiting dilution with an expected cell number of either 0.5 or 2 per well in 96-well plates with 3 replicate plates per condition and cell number. Cloning efficiency was calculated as the number of wells with colonies divided by the number of expected colonies based on the number of cells plated. (D) Cloning efficiency for *ATRX* knockouts in *TP53* knockout genetic background.

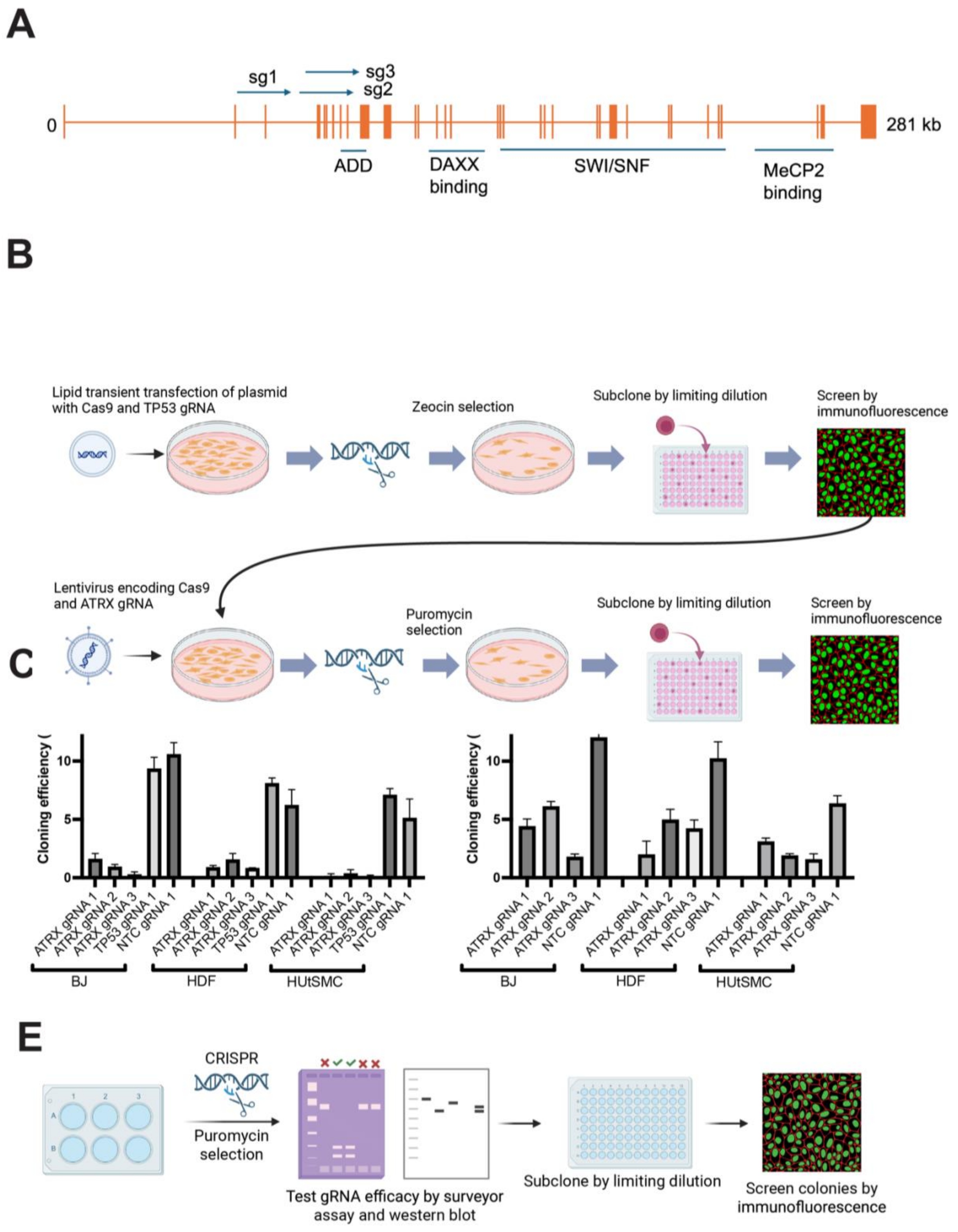
A
sg1
sg3
sg2
0
281 kb
ADD
DAXX binding
SWI/SNF
MeCP2 binding
B
Lipid transient transfection of plasmid with Cas9 and TP53 gRNA
Zeocin selection
Subclone by limiting dilution
Screen by immunofluorescence
Lentivirus encoding Cas9 and ATRX gRNA
Puromycin selection
Subclone by limiting dilution
Screen by immunofluorescence
C
Cloning efficiency (
10
5
0
ATRX gRNA 1
ATRX gRNA 2
ATRX gRNA 3
TP53 gRNA 1
NTC gRNA 1
BJ
HDF
HUtSMC
Cloning efficiency (
10
5
0
ATRX gRNA 1
ATRX gRNA 2
ATRX gRNA 3
NTC gRNA 1
BJ
HDF
HUtSMC
E
CRISPR
Puromycin selection
Test gRNA efficacy by surveyor assay and western blot
Subclone by limiting dilution
Screen colonies by immunofluorescence

**Supplemental Figure S6. Co-occurrence of *ATRX* mutation with *TP53* and *RB1* alterations.**

Oncoprint plots demonstrating co-occurrence of *ATRX* alterations with *TP53* and *RB1* alterations in TCGA (A) and GENIE (B) sarcoma cohorts.

A

TCGA sarcoma

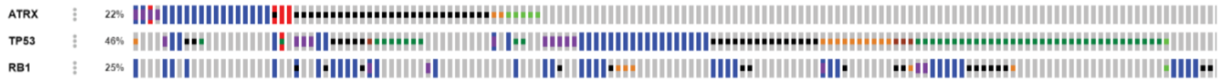


| A | B | Neither | A Not B | B Not A | Both | Log2 Odds Ratio | p-Value | q-Value ▲ | Tendency |
|---|---|---|---|---|---|---|---|---|---|
| TP53 | RB1 | 128 | 62 | 8 | 55 | >3 | <0.001 | **<0.001** | **Co-occurrence** |
| ATRX | TP53 | 111 | 25 | 86 | 31 | 0.678 | 0.131 | 0.164 | Co-occurrence |
| ATRX | RB1 | 152 | 38 | 45 | 18 | 0.678 | 0.164 | 0.164 | Co-occurrence |

B

GENIE sarcoma

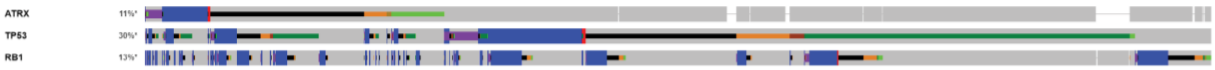


| A | B | Neither | A Not B | B Not A | Both | Log2 Odds Ratio | p-Value | q-Value ▲ | Tendency |
|---|---|---|---|---|---|---|---|---|---|
| TP53 | RB1 | 3144 | 923 | 161 | 564 | >3 | <0.001 | **<0.001** | **Co-occurrence** |
| ATRX | TP53 | 3132 | 173 | 1120 | 367 | 2.569 | <0.001 | **<0.001** | **Co-occurrence** |
| ATRX | RB1 | 3753 | 314 | 499 | 226 | 2.436 | <0.001 | **<0.001** | **Co-occurrence** |

**Supplemental Figure S7. Validation of p53 knockout.**

(A-B) Western blotting for TP53 in BJ and HDF fibroblasts (A) and HUtSMC (B). The red arrow indicates p53; the lower molecular weight band is non-specific. (C) To confirm p53 functional status, we blotted for a downstream p53 effector, p21, after exposure to the DNA damaging agent doxorubicin.

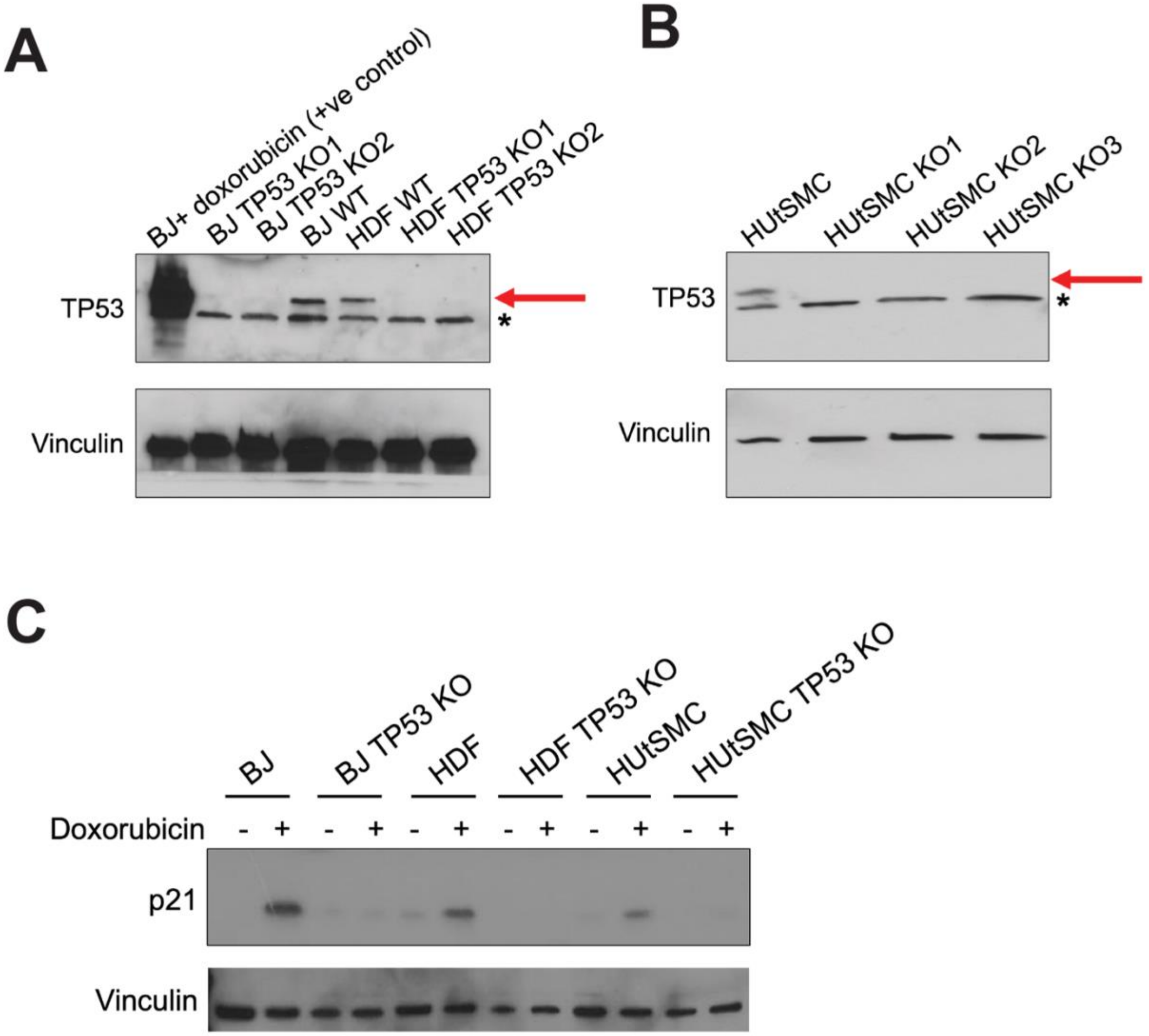

**Supplemental Figure S8. ATRX loss is associated with genomic instability.**

(A) Representative histograms from flow cytometry for cell cycle analysis after propidium iodide staining. (B) Fraction of the genome altered in *ATRX* mutant versus WT sarcomas in TCGA. (C) Aneuploidy score in *ATRX* mutant versus WT sarcomas in TCGA. (D) TERT mRNA expression in *ATRX* mutant versus WT sarcomas in TCGA. Plots in B-D made in cBioPortal.

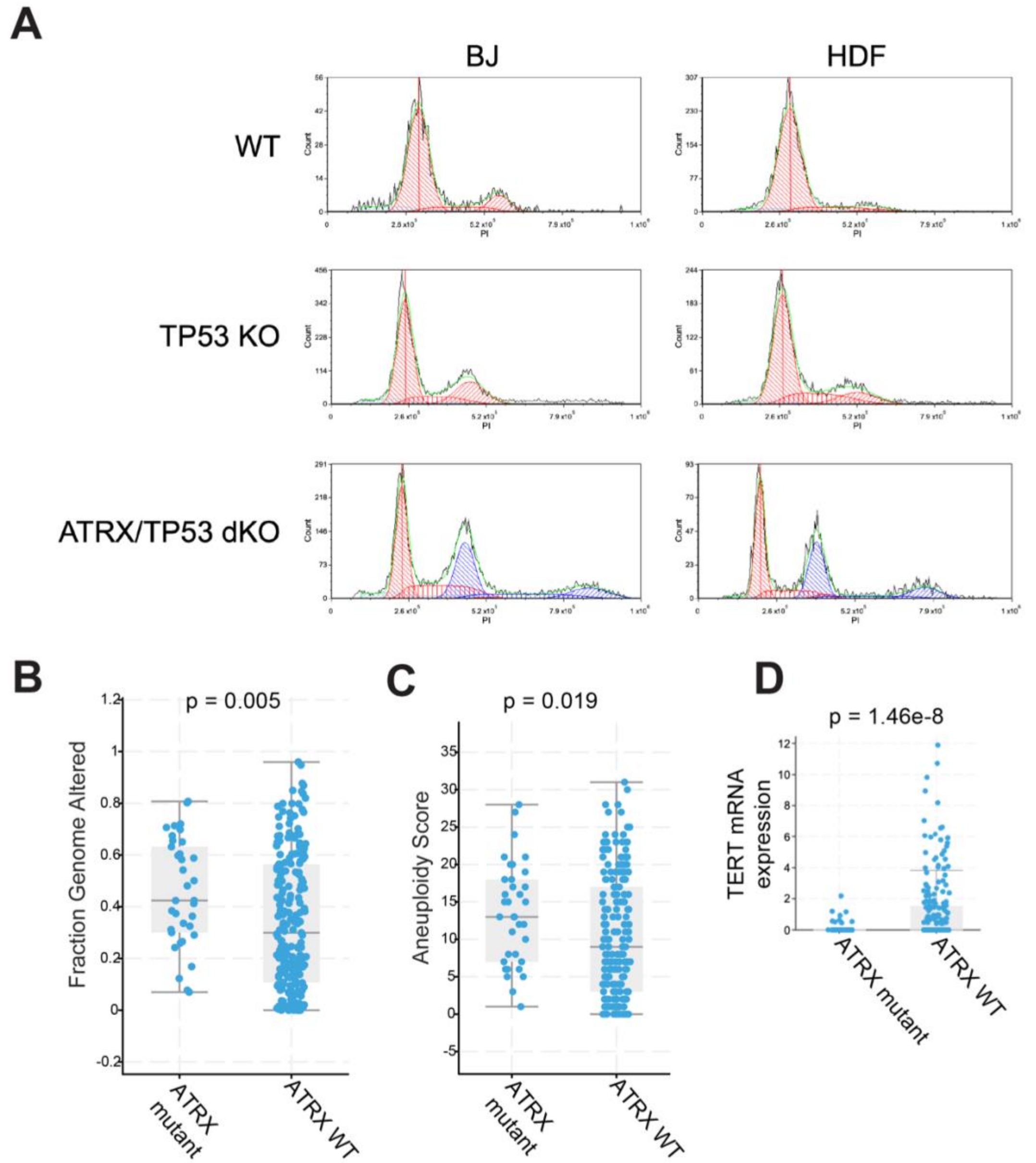

**Supplemental Figure S9. ATRX loss in human UPS is not associated with ALT.**

(A) Representative images of TEL-FISH showing ALT positive (top) versus negative (bottom) tumors. Scale bars = 50 μm. (B) Percent of tumors with ALT presence or negativity. (C) Percent of ALT positivity based on ATRX expression by IHC.

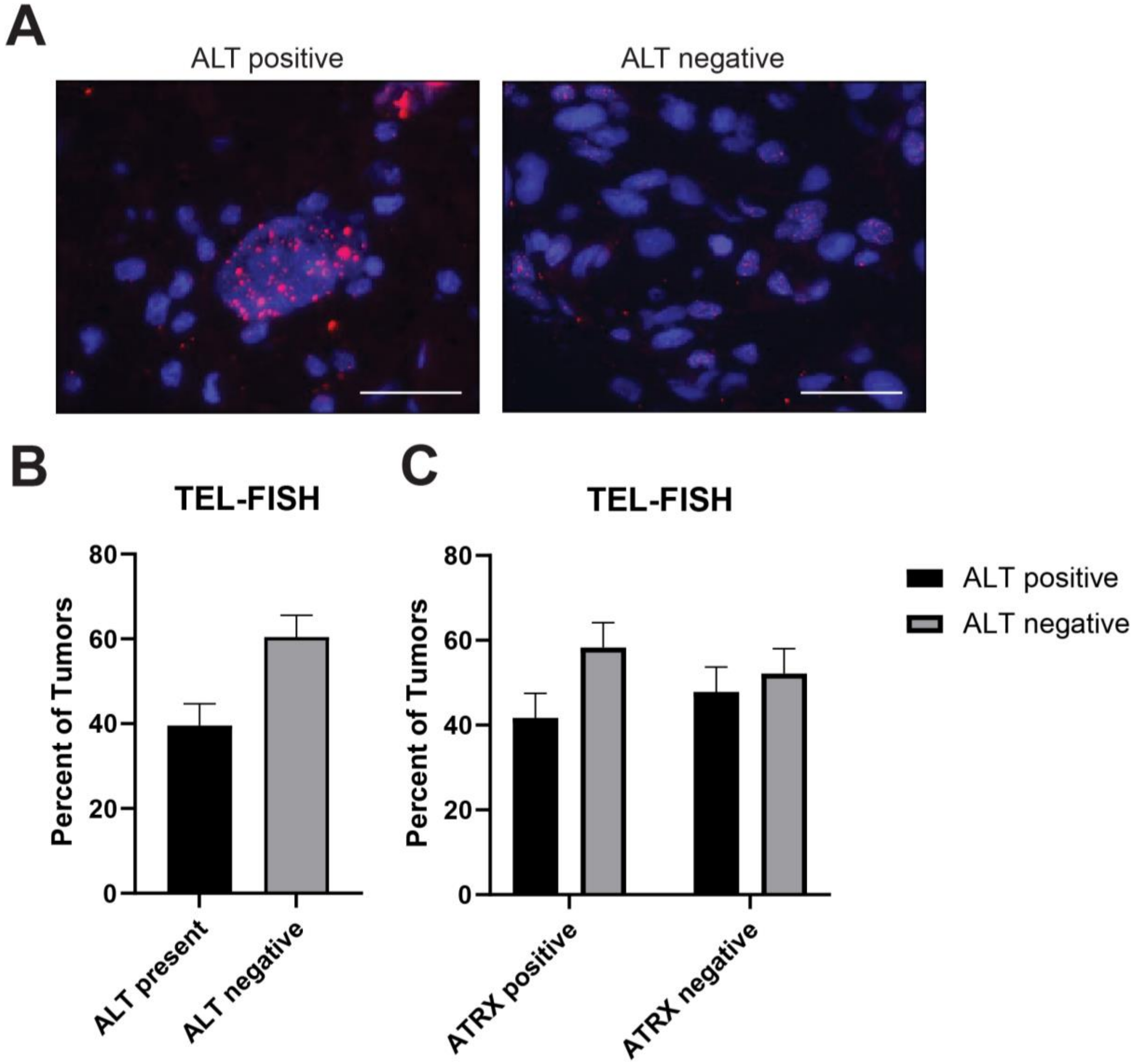

**Supplemental Figure S10. Depletion of ATRX with CRISPR in UPS cell lines.**

(A) Western blot of ATRX in untreated cell lines. UPS186, UPS271.1, RIS620, RIS819.1, UPS060, and UPS511 are patient-derived UPS cell lines. LN18 is a glioblastoma cell line, hMSC is human mesenchymal stem cells, and these served as positive controls. (B) Western blotting for ATRX in polyclonal populations of UPS cell lines following expression of 3 different all-in-one CRISPR plasmids. Three different gRNAs were screened. P indicates parental control (untreated cells).

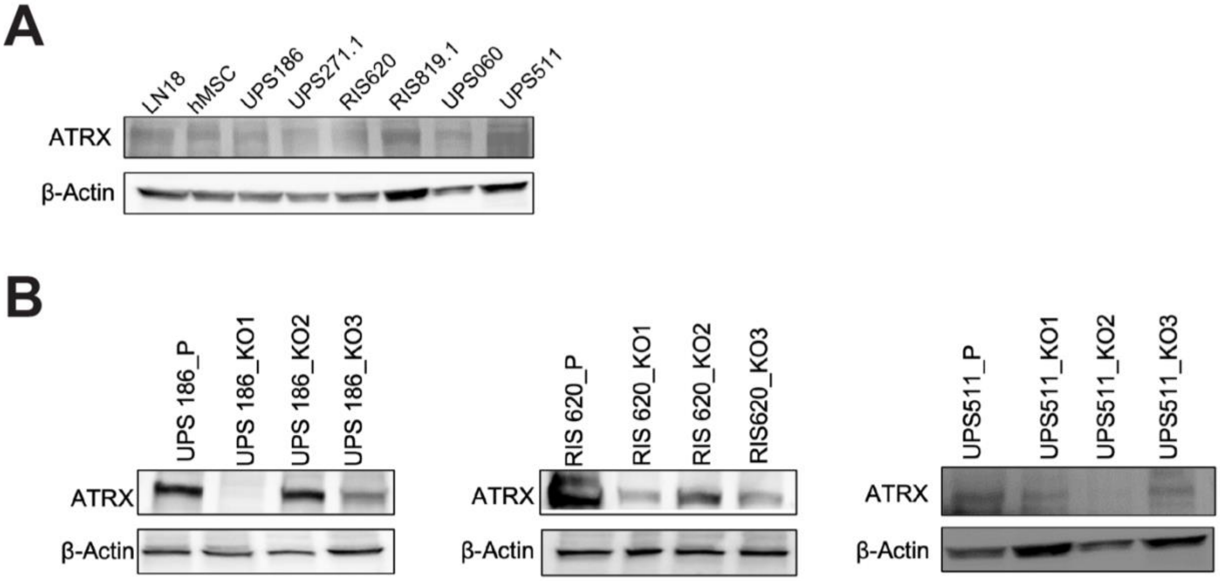

**Supplemental Figure S11. Upregulated motifs in ATRX KO cells.**

(A-B) IGV snapshot showing ATAC-seq signal tracks for genomic locus harboring ZNF404 (A) and GALNT2 (B) genes, 2 of the most upregulated genes in *ATRX* KO cells. (C) UMAP analysis utilizing ATAC-seq data. ATRX KO conditions are colored in dark purple and *ATRX* WT conditions are colored in light purple.

Supplemental Figure 11 . ATAC-seq

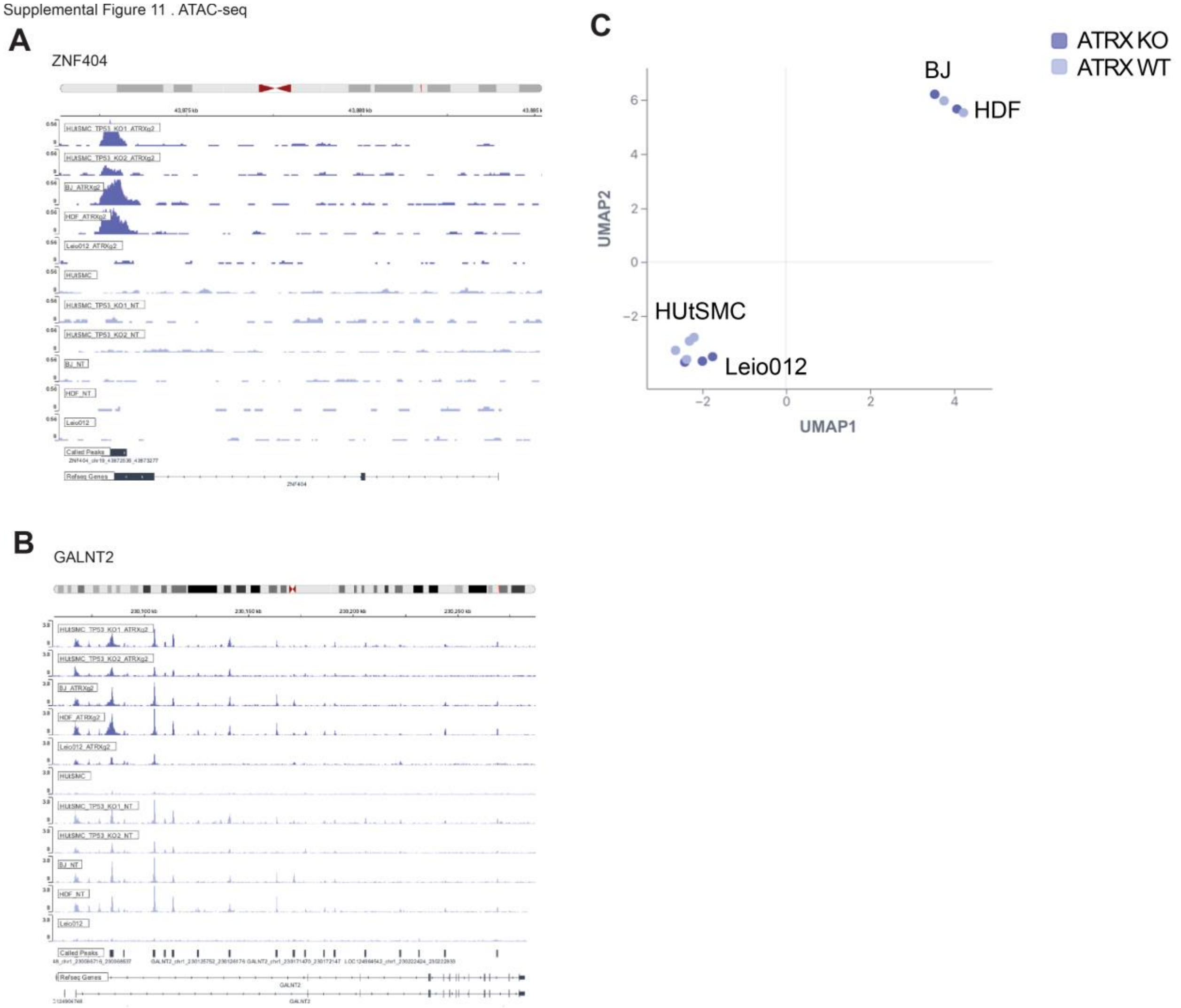

**Supplemental Figure S12. Alterations in *PRDM4* and *NFIX* in sarcoma TCGA.**

(A-B) PRDM4 genomic alterations (A) and combined genomic and transcriptomic alterations (B) by cancer subtype in TCGA. The red arrow highlights the sarcoma dataset. (C-E) Overall survival (C), disease-specific survival (D), and progression-free survival (E) based on the presence or absence of PRDM4 overexpression (OE). (F-G) NFIX genomic alterations (F) and combined genomic and transcriptomic alterations (G) by cancer subtype in TCGA. The red arrow highlights the sarcoma dataset. (H-J) Overall survival (H), disease-specific survival (I), and progression-free survival (J) based on the presence or absence of NFIX overexpression (OE). (K-N) Gene expression of the 4 NFI genes based on ATRX expression. NS = not significant. P values on Kaplan Meier curves are from log-rank tests.

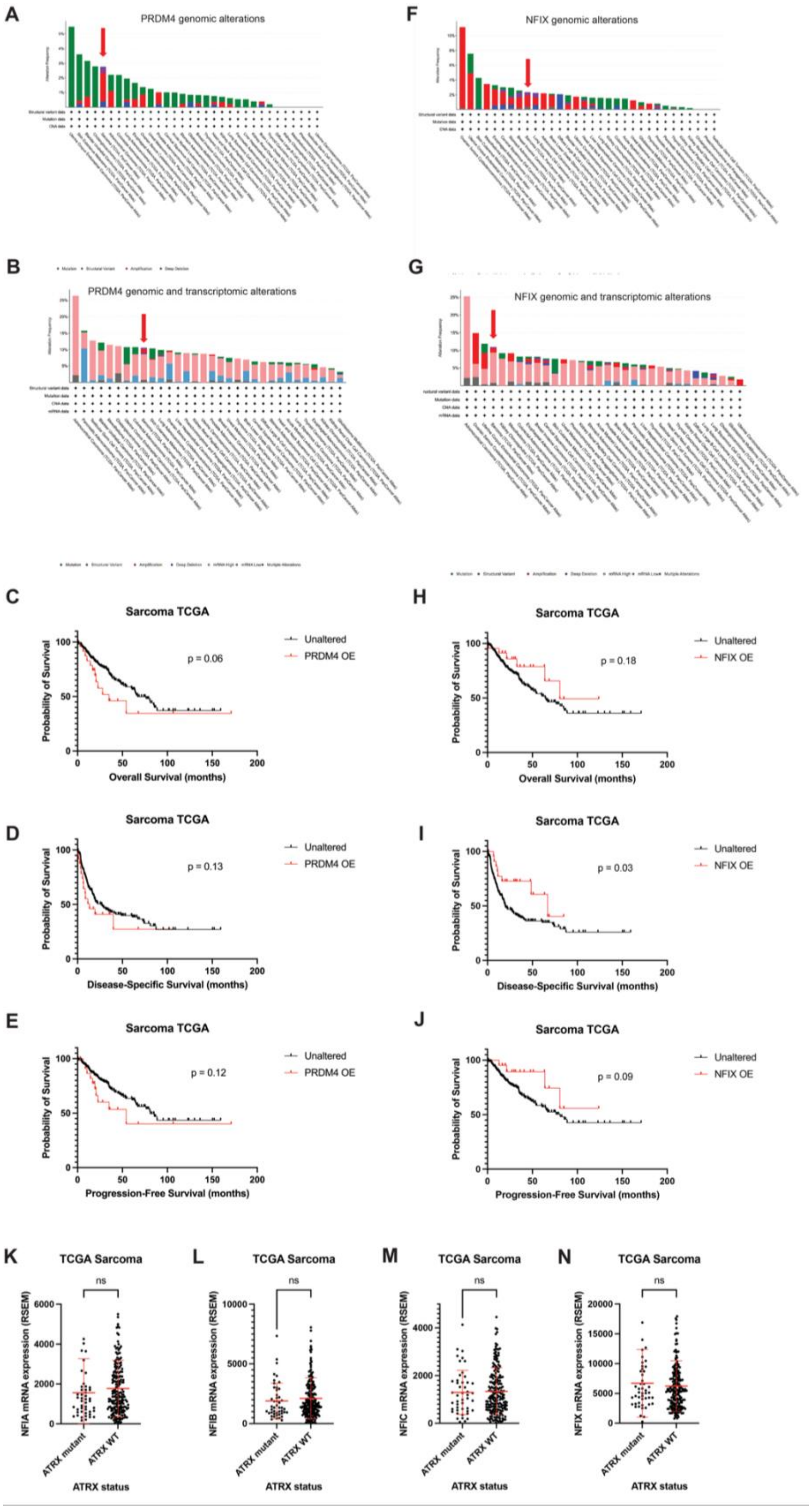
A
PRDM4 genomic alterations
F
NFIX genomic alterations
B
PRDM4 genomic and transcriptomic alterations
G
NFIX genomic and transcriptomic alterations
C
Sarcoma TCGA
Unaltered
PRDM4 OE
p = 0.06
Probability of Survival
Overall Survival (months)
D
Sarcoma TCGA
Unaltered
PRDM4 OE
p = 0.13
Disease-Specific Survival (months)
E
Sarcoma TCGA
Unaltered
PRDM4 OE
p = 0.12
Progression-Free Survival (months)
H
Sarcoma TCGA
Unaltered
NFIX OE
p = 0.18
Overall Survival (months)
I
Sarcoma TCGA
Unaltered
NFIX OE
p = 0.03
Disease-Specific Survival (months)
J
Sarcoma TCGA
Unaltered
NFIX OE
p = 0.09
Progression-Free Survival (months)
K
TCGA Sarcoma
ns
NFIA mRNA expression (RSEM)
ATRX mutant
ATRX WT
ATRX status
L
TCGA Sarcoma
ns
NFIB mRNA expression (RSEM)
ATRX status
M
TCGA Sarcoma
ns
NFIC mRNA expression (RSEM)
ATRX status
N
TCGA Sarcoma
ns
NFIX mRNA expression (RSEM)
ATRX status

**Supplemental Figure S13. Correlation with ITGB2.**

(A) Correlation of *ITGB2* with *PRDM4* in sarcoma TCGA cohort. (B) *ITGB2* expression in *ATRX* mutant versus WT in TCGA sarcoma database.

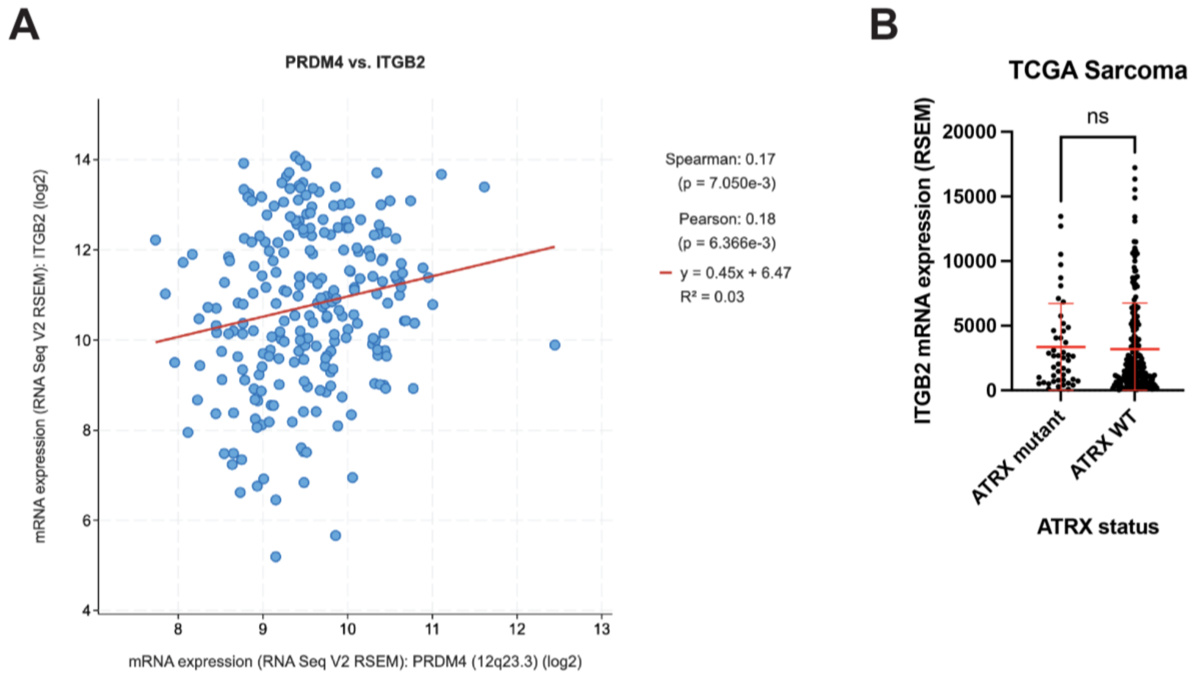

**Supplemental Figure S14. Analysis of PRMT5 targets.**

PRDM4 interacts with PRMT5. Western blots showing PRMT5 products in the indicated BJ and HDF genotypes. Nuclear extracts were used for H3R8me2s and H4R3me2s blots. whole cell extracts were used for SDMA blot.

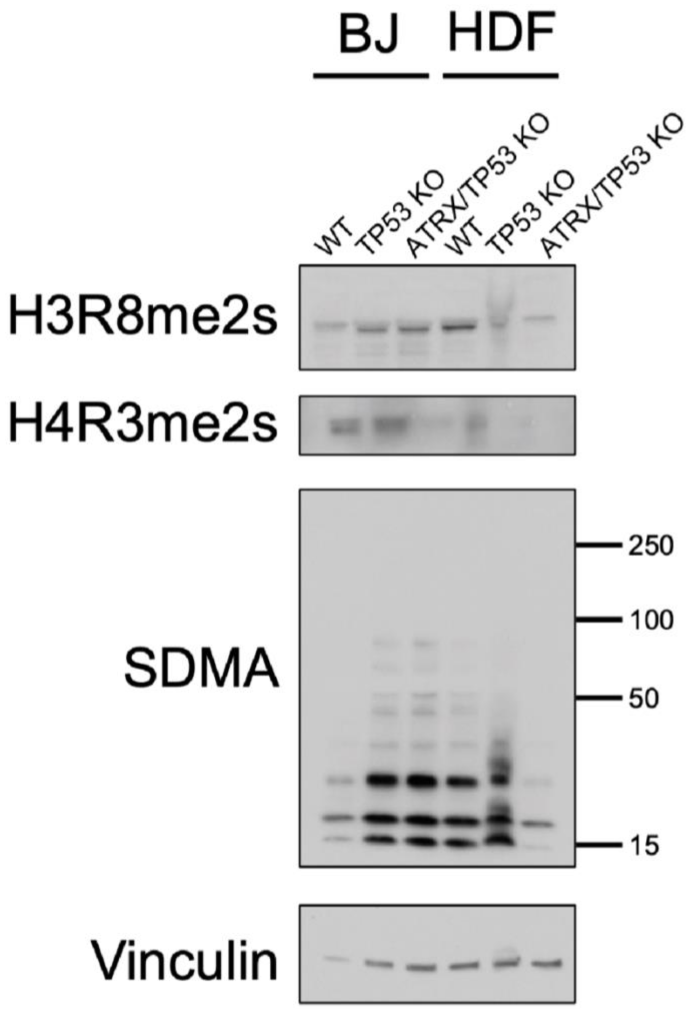

**Supplemental Figure S15. Identification of ATRX knockout cell lines.**

Western blotting for ATRX in a panel of MPNST cell lines (A) and DDLPS cell lines (B).

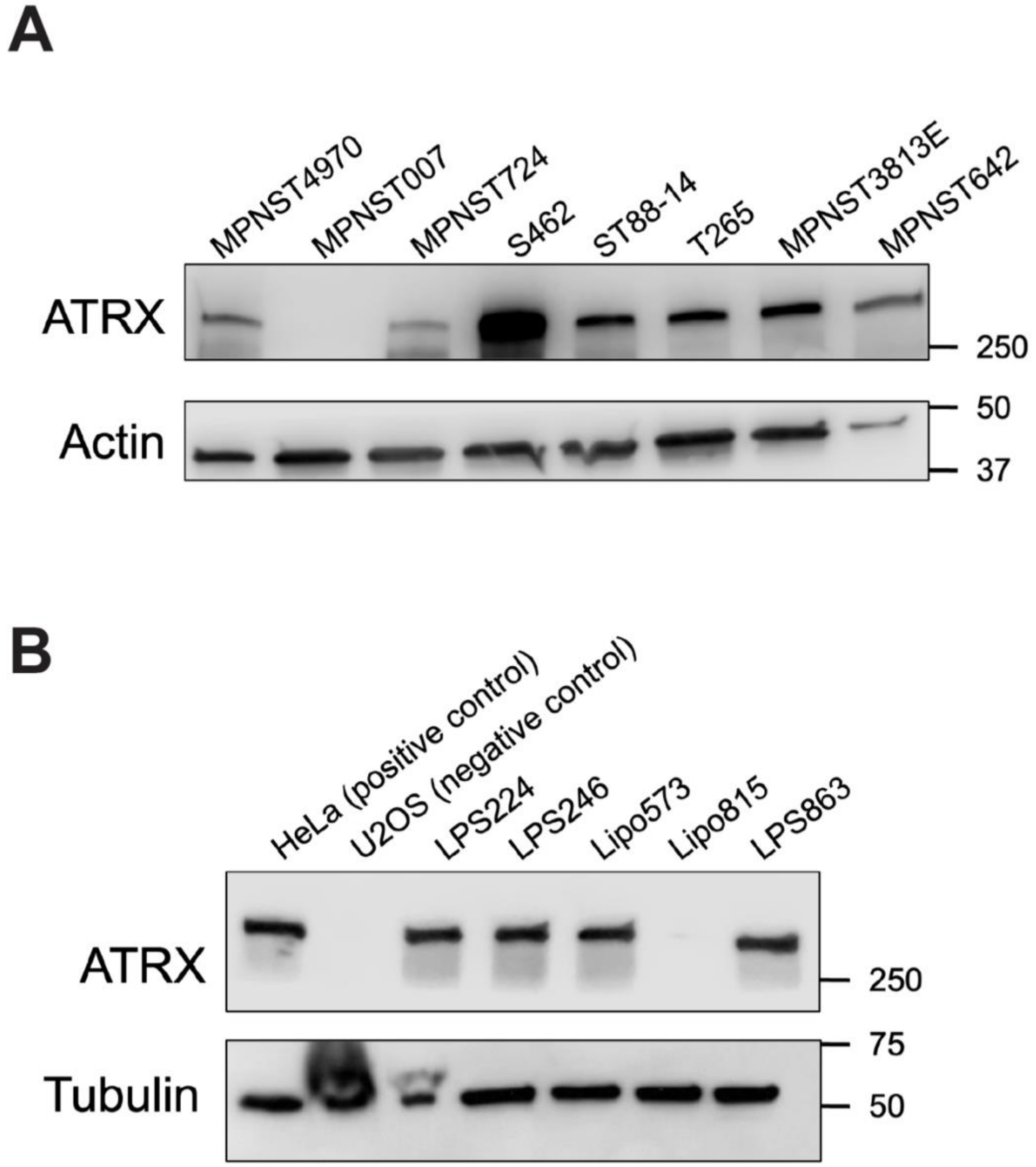

**Supplemental Figure 16. Validation of PRDM4 and NFIX antibodies.**

(A) Nuclei were isolated from Lipo815 and MPNST007 cell lines, and nuclear and cytoplasmic extracts were probed for PRDM4 and NFIX to validate that these antibodies recognize nuclear proteins. (B) Immunofluorescence staining for PRDM4 and NFIX, demonstrating nuclear localization. Scale bar = 50 μm.

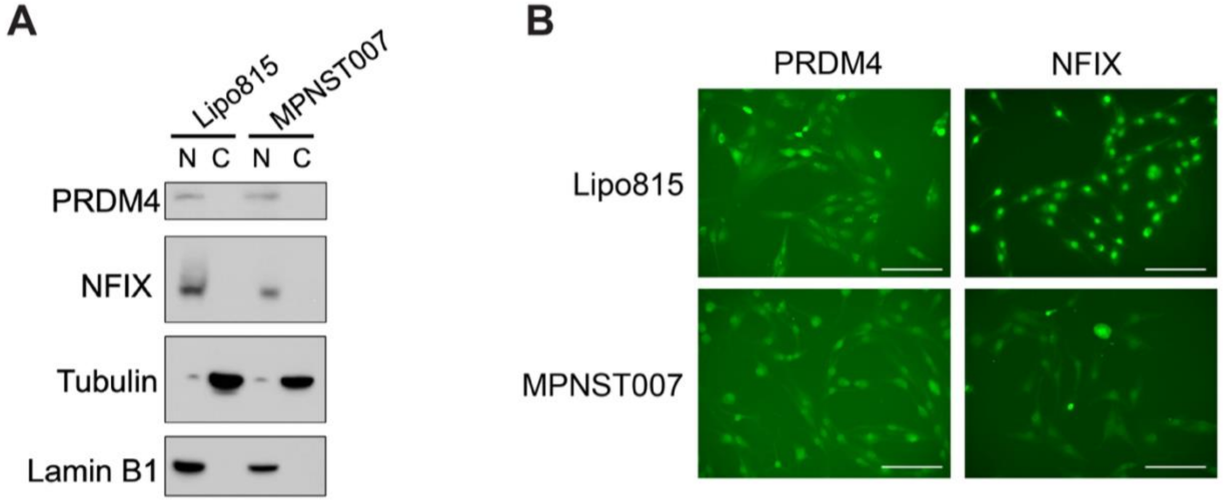

**Supplemental Figure S17. YAP1 levels do not change significantly with ATRX loss.**

YAP1 western blotting in BJ, HDF, and HUtSMC cells.

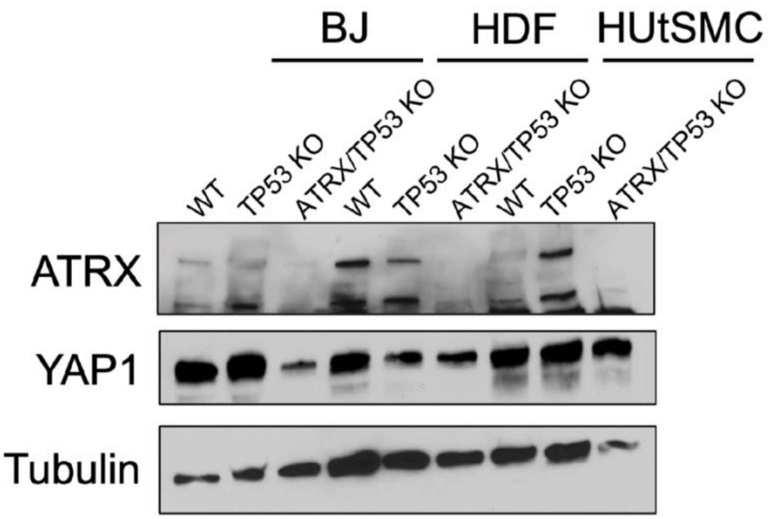
